# Roquin exhibits opposing effects on RNA stem-loop stability through its two ROQ domain binding sites

**DOI:** 10.1101/2024.11.22.624803

**Authors:** Jan-Niklas Tants, Andreas Walbrun, Lucas Kollwitz, Katharina Friedrich, Matthias Rief, Andreas Schlundt

## Abstract

The interaction of mRNA and regulatory proteins is critical for post-transcriptional control. For proper function, these interactions as well as the involved protein and RNA structures are highly dynamic and thus, mechanistic insights from structural biology are challenging to obtain. In this study, we employ a multifaceted approach combining single-molecule force spectroscopy with NMR spectroscopy to analyze the concerted interaction of the two RNA-binding interfaces (A-site, B-site) of the immunoregulatory protein Roquin’s ROQ domain with the 3’ untranslated region (UTR) of the *Ox40* mRNA. This 3’UTR contains two specific hairpin structures termed constitutive and alternative decay elements (CDE, ADE), which mediate mRNA degradation via binding of Roquin. Our single-molecule experiments reveal the CDE folds cooperatively, while ADE folding involves at least 3 on-pathway and 3 off-pathway intermediates. Utilizing an integrated microfluidics setup allows to extract binding kinetics to Roquin in real time. Supported by NMR, we find opposing effects of the two Roquin sub-domains on distinct regions of the ADE: while the A-site interacts strongly with the folded apical stem-loop, we find that the B-site has a distinct destabilizing effect on the central stem of the ADE owed to single-strand RNA binding. We propose that RNA-motif nature and Roquin A- and B-sites jointly steer mRNA decay with context-encoded specificity, and we suggest plasticity of stem structures as key determinant for Roquin-RNA complex formation. The unique methodological combination of NMR and single-molecule force spectroscopy reveals an unknown mechanism of a dual-function RNA-binding domain suggesting a new model for target RNA recognition.

**SIGNIFICANCE STATEMENT:** Local RNA structure is decisive for specific engagement with gene-regulatory proteins and, as a consequence, correct cellular function. However, its existence often appears dynamic and thus, challenging to study. This study shows how NMR and single-molecule force spectroscopy efficiently complement each other to provide high-resolution, time-resolved data on RNA folding intermediates during dynamic complex formation with the immune-regulating protein Roquin, which exploits multiple RNA-binding sites. Our data reveal a dual-mode binding of Roquin to RNA by firmly attaching to the stem-loop and, at the same time, destabilizing other regions making them accessible to downstream interaction partners.

## INTRODUCTION

RNA *cis* elements contribute to the regulation of key cellular processes, e.g. translational control. Often, cognate *trans*-acting factors, i.e. proteins, confer functionality to the underlying RNA-protein complexes. High specificity in RNA target recognition by proteins is key for a functional outcome, and both proteins and RNAs contribute to this(1). Proteins often achieve specificity in ribonucleoprotein (RNP) formation via their modular architecture or specialized RNA-binding domains, e.g. dsRBDs for double-stranded RNAs. Despite their limited tertiary contacts, RNAs exhibit a great structural variety. Due to inherent flexibility, RNA *cis* elements sample a large conformational space and thus, one sequence can provide binding interfaces for more than one protein(2). Ultimately, the spatio-temporal availability of a certain RNA structure will govern final *cis*-*trans* pair formation and affect downstream functions, which remains extremely difficult to predict, and often, not straightforward to analyze on a structural level(1).

The immune-regulatory protein Roquin is specialized for RNA hairpin recognition and mediates mRNA decay through *cis* element binding(3). The A-site located in its coreROQ domain interacts with single-stranded (ss) loops through base-specific contacts, while backbone contacts to apical stems assure shape recognition(4, 5). Additionally, structured extensions flanking the coreROQ domain and constituting the extended ROQ domain (extROQ) were suggested to interact with double-stranded RNAs through a B-site(6). Up to date it remains elusive whether Roquin interacts with both types of RNAs simultaneously and whether both RNA structures are recognized within one *cis* element. An apparent increase in affinity by binding of both Roquin A- and B-sites suggested complex stabilization via the B-site binding to duplex regions distal to the hairpin(7). Roquin recognizes two types of hairpins, which contain a tri- or a hexaloop, named constitutive and alternative decay element (CDE(4) and ADE(5), respectively) and were shown to be functionally redundant in the *Ox40* 3’UTR. Despite that both *cis* elements exploit the same Roquin binding interface, their overall shapes differ significantly(7, 8). While CDEs and ADEs are usually stable motifs(2, 7), Roquin does not require preformed stem-loop structures for initial engagement(2). Its capability to induce secondary structure and the structural variety of CDEs and ADEs challenge a simple model of combined recognition of hairpin and double-stranded regions through the Roquin A- and B-sites(8). Previous studies exploited only small isolated elements or a full 3’UTR context, which hampers sufficient probe coverage of individual *cis* elements due to its size. Not only for Roquin, we thus currently lack detailed structural insights in the concerted interaction of both protein interfaces with target RNAs and a composite understanding of target specificity and context-related mRNA suppression.

As the example of Roquin shows, studying the structure and dynamics of large complexes of RNA and proteins generally poses a challenge for structural biology and often a single technique is not able to provide a complete picture. While NMR (nuclear magnetic resonance) can provide detailed structural and kinetic information at the atomic level, larger structures are often elusive or not sufficiently accessible to NMR with the required resolution. As a consequence, detection and quantification of intermediates, especially in RNAs, remains challenging. Single-molecule force spectroscopy (SMFS) has given detailed insight into the dynamics of proteins and nucleic acids(9–11) ranging from microseconds to the timescale of minutes. A strength of SMFS is that it can be applied to large systems and still provide structural information on folding pathways and associated intermediates, at least on the primary structure level(12), by steering the molecules through their folding free-energy landscapes. In this, even rare events populated to less than 0.01% can be detected(13). So far, studies on RNA structures have predominantly investigated dynamics and transition path times of folding(14–17). However, the interaction with cognate RNA-binding proteins, as is integral for Roquin-binding *cis*-elements, adds an additional layer of complexity to RNA dynamics and stability, a problem we sought to address in this study.

We here combine the structural resolution provided by NMR with the ability of optical tweezers (OT) as SMFS technique to observe the formation of complexes between RNA and proteins in real time. OT experiments confirm the overall fold of the full-length (fl) *Ox40* 3’UTR at nucleotide resolution (**Fig. 1A**), based on which we provide a detailed folding landscape of the extended *Ox40* ADE. While we observe stabilization of apical hairpins through the Roquin A-site, we give evidence for unprecedented ssRNA binding through the B-site, leading to a destabilization of basal stem regions, using an interdisciplinary approach supported by bulk biochemical assays such as electrophoretic mobility shift assays (EMSA), CD spectroscopy and hydroxyl radical probing. We precisely quantify conformational populations of ADE folding intermediates and observe a Roquin-induced shift in folding equilibrium. Our data shows that Roquin contributes to the structural plasticity of RNAs selectively exploiting its two RNA-binding sites. Binding of Roquin to ssRNAs partially explains the modulatory effect on RNA structure and further expands the known RNA target spectrum. This study serves a paradigm on how SMFS and NMR spectroscopy supplemented with biophysical techniques provide time-resolved and quantifiable structure snapshots of RNA folding intermediates, important for and applicable to dynamic RNPs.

**Figure 1.**
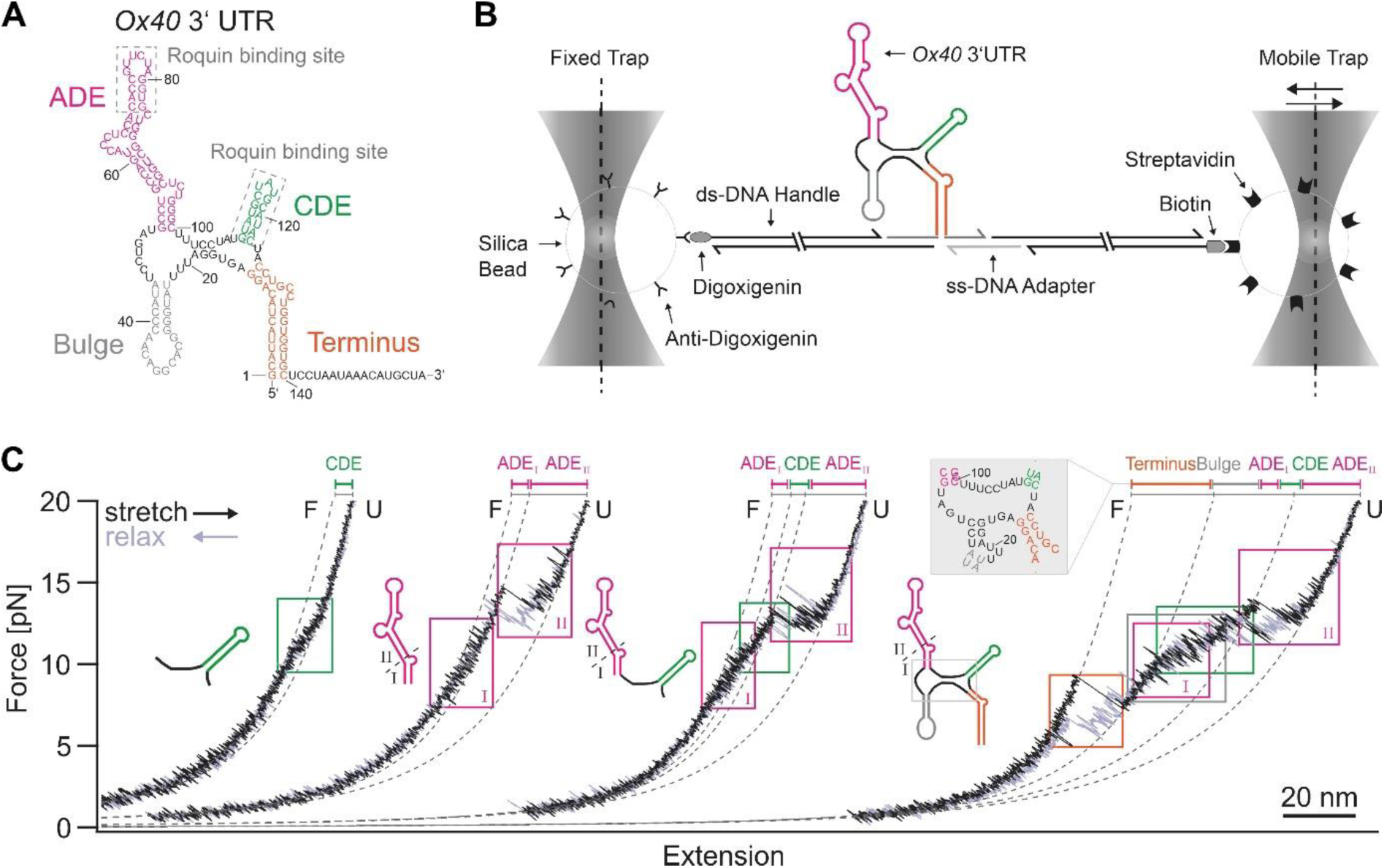
OT measurements capture the modular architecture of the *Ox40* 3’UTR. **A)** Secondary structure of the fl *Ox40* 3’UTR as determined in (7). Individual structured/functional elements are color-coded. **B)** Setup and measurement principle of the dual-beam OT system used in this study (for details, see SI). **C)** Representative force-extension curves of the isolated CDE, ADE, the combined ADE-CDE construct and the fl 3’UTR construct. Note that the extension axis does not display absolute values since the traces are arranged for side-by-side comparison. Unfolding/refolding transitions of structural elements are highlighted with boxes, and WLC fits to the most pronounced intermediates are shown as dashed lines. The unfolded contour length contribution of the Terminus obtained in the fl force-extension curves reveals an additional stemmed unit adjacent to the Bulge (gray inset). In **Fig. S2**, we show a collection of five additional sample traces for each of the constructs used.

## RESULTS

### OT experiments capture structure and folding intermediates of the fl *Ox40* 3 UTR

We initially used an OT setup to study folding and unfolding of RNA structure within the *Ox40* 3’UTR. To this end, we designed a dumbbell construct where a single RNA molecule is tethered between two beads as shown in **Fig. 1B** (see Material and Methods and (18) for more details). We investigated four different constructs with increasing complexity(7): the isolated CDE, the isolated ADE, a tandem ADE-CDE construct, and the fl *Ox40* 3’ UTR (**Fig. 1C**).

**Fig. 1C** shows representative examples of stretch (black) and relax (gray) cycles for each construct. The trace of the isolated CDE showed a rapid two state folding/unfolding transition at 11.5 pN. Note that only the hump-like transition visible in the green box reflects CDE folding and unfolding, while the general convex shape of the trace is owed to the entropic elasticity of the DNA handles(19). Fits using the worm-like chain (WLC) model of polymer elasticity(20, 21) allow precise determination of the contour length gain during an unfolding event. We find an average length gain of 7.8 ± 0.2 nm (**Table S1**) in agreement with the expected value of 15 bases transitioning from the folded to the fully extended state. While the smoothed trace filters out individual transitions, higher-bandwidth data in **Fig. S1 A** and **B** allow direct observation of folding/unfolding fluctuations happening on the sub-ms timescale (**Fig. S1 C and D**).

For the isolated ADE, we find two prominent folding/unfolding transitions (**Fig. 1C**). A first transition (I) occurs at around 9 pN, followed by a more prominent transition (II) at 15 pN. Transition I is consistent with unfolding of the basal stem until the first interior bulge G56-C93, while transition II subsequently involves full unfolding of the remainder of the ADE. The length gains obtained by WLC fits (see **Table S1**) are in full agreement with this interpretation.

Unfolding/refolding of the combined ADE-CDE construct exhibits features of both isolated elements and is consistent with the structure shown in **Fig. 1A**. At low forces, we find transition I of ADE followed by the hump characteristic for CDE unfolding and finally, transition II of ADE unfolding as the last and most stable structural component. Again, the contour length gains determined by WLC fits match this interpretation (**Table S1**). We find no indication of additional tertiary structural interactions between the two elements, which supports earlier findings of the two elements to exist independently from each other(7).

The fl *Ox40* 3’UTR construct additionally contains two new signatures that represent unfolding of the Terminus and the Bulge. Based on the secondary structure predicted by *mfold*(22) (**Fig. 1A**), one would expect the unfolding of the Terminus to either stop at the GGA bridge (nts 17-19) or to unfold even further in the same step, leaving only the CDE, ADE and Bulge elements closed. Interestingly, the contour length gain extracted from the WLC fit (**Table S1**) after the first unfolding transition (orange box) matches neither of these assumptions. This finding suggests that the GGA nucleotides (17–19) rather pair with UCC adjacently to the Bulge corresponding to the core region structure shown in the close-up of **Fig. 1C**. This structure is in line with the lowest free-energy structure predicted by *Vienna RNAfold*(23) **(Fig. S1E**)(7), demonstrating the capability of SMFS for tackling ambiguities between RNA structure prediction tools.

Unfolding of the Terminus opens the structure such that all the remaining folded stem-loops in the structure are subject to force in parallel. Consequently, this leads to simultaneous folding/unfolding transitions of the remaining structural parts. We therefore find their individual signatures overlapping and strict separation of the individual transitions becomes impossible with the exception of the ADE_II_ transition, which, owing to higher mechanical and energetic stability(7), occurs lastly. Importantly, no additional folding event occurs when comparing individual elements with the fl 3’UTR. Notably, we found that after complete unfolding, the fl *Ox40* 3’UTR can be trapped in a misfolded state after the chain is relaxed to zero force which is likely mediated by a non-native fold of the Terminus (**Fig. S2**). Such behavior is well known for RNA constructs of similar complexity(24, 25) and thus underlines the critical role of controlled mRNA folding in its native context. Altogether, our OT experiments confirm the structural independence of the four elements found in the *Ox40* 3’UTR and disclose its total folding/unfolding network to largely result from the autonomous folding/unfolding of those elements. This qualifies the OT methodology for utilization in detailed interrogation of RNA-structural contexts.

### The *Ox40* ADE samples multiple folding intermediates

We have previously shown that a CDE hairpin structure can exchange between a folded and unfolded state, both of which recognized by individual RBPs(2). We thus asked whether a yet more pronounced intrinsic heterogeneity – affected by Roquin – is observed in the *Ox40* ADE, providing a more complex fold than the CDE.

We first set out to determine the stability of individual ADE segments in the apo state and to capture its full conformational space. The high melting point of ∼75 °C of the ADE described before(7) suggests formation of a network of stabilizing H-bonds. We recorded HNN-COSY(26) NMR spectra to detect and quantify basepair-contained H-bonds (**Fig. 2A**). Indeed we detected H-bonds for most basepairs described before(7), confirming formation of strong duplex regions (**Fig. 2A**) interspersed with bulges. Increased temperatures led to destabilization of the majority of basepairs (**Fig. S3**), as expected. However, the apical stem and the duplex region below the central bulge (residues 61-67) showed detectable imino proton signals at 37 °C, indicative of secondary structure. Base pairs of residues 67-70 with residues 83 and 85-87 were not directly observed likely owed to the lower stability of this region. C84 and the adjacent central bulge render this stretch less stable, as indicated by the absence of imino proton signals (**Fig. 2A**). In sum, the NMR data support the outlined secondary structure scheme of the *Ox40* ADE on the level of individual basepair stabilities.

**Figure 2.**
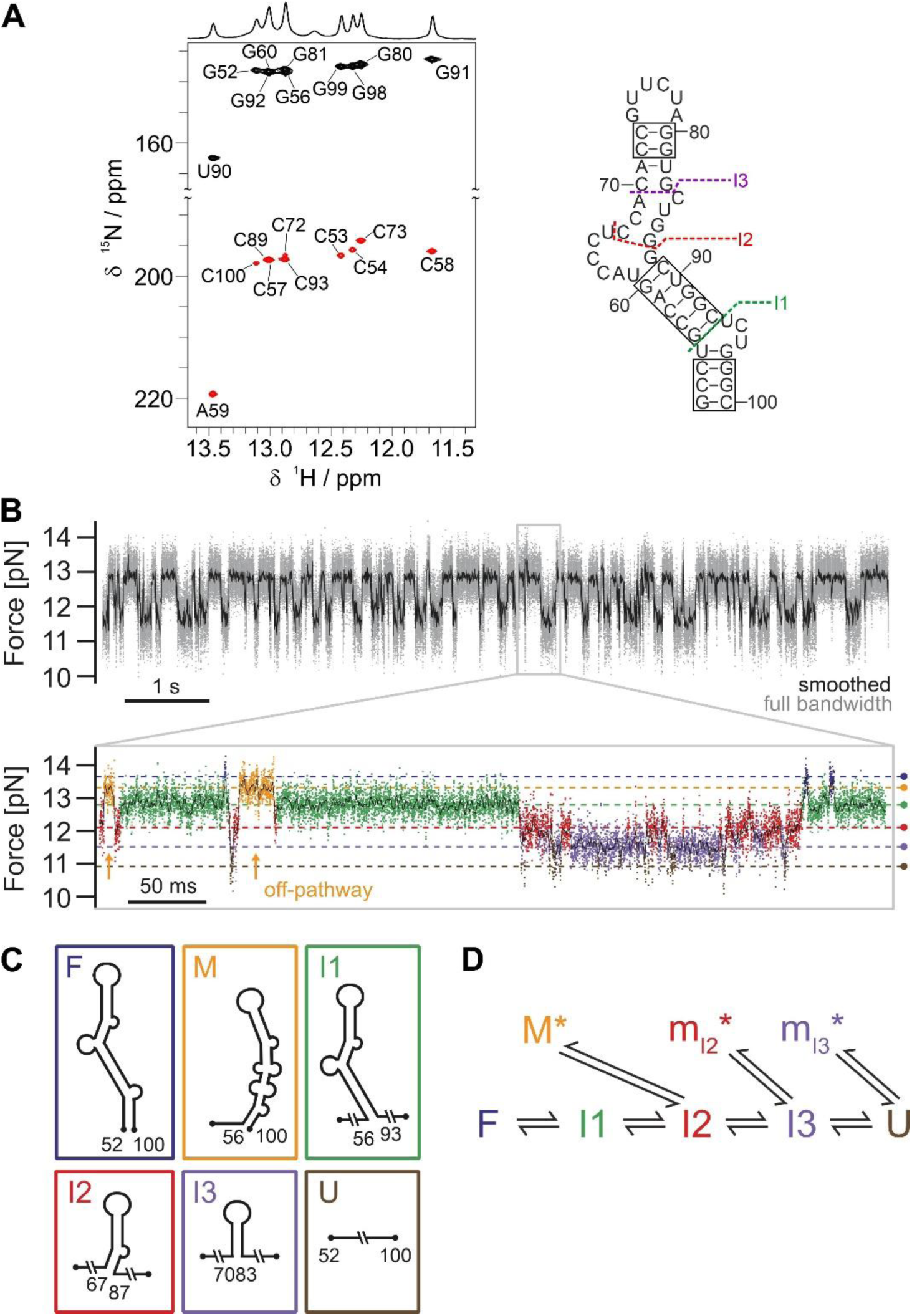
The *Ox40* ADE adopts a stable fold via multiple folding intermediates. **A)** HNN-COSY experiment of apo ADE. H-bond donor peaks are shown in black, pairing H-bond acceptor peaks in red. Observed H-bonds confirm base pairs, as indicated in boxes on the ADE secondary structure on the right. Dotted lines show the position of the intermediates extracted from OT experiments below corresponding to structures shown in D). **B)** Passive-mode measurement of the ADE. States are more folded towards higher forces and more unfolded at lower forces. Note that the folding process shortens the length of the construct, and hence, the beads are pulled out of the trap, increasing the load. The zoom into a 450 ms excerpt of this trace reveals six different states constituting the folding network, including three intermediates (green, red, and violet) and a misfolded state (orange). Appearances of the off-pathway misfold (orange) are highlighted with arrows to emphasize their exclusive accessibility from the red state. **C)** Schematic depiction of the states forming the unfolding/refolding network of the ADE (see color code from B)). The misfolded orange state is formed by non-native folding downstream of I2. The schematic structure M represents the most likely secondary structure based on its number of unfolded nucleotides, supported by predictions of the *RNAstructure fold* server(44) (see **Fig. S4B** for more details). **D)** Folding network of the ADE extracted from transitions observable in passive mode. While lifetimes of F, I1 and U are single-exponentially distributed, we see multi-exponential distributions on the force levels of the purple, red and orange states (**Fig. S4**, ADE pathway), indicating ensembles of states with short-lived misfolds (indicated by an asterisk).

We next used OT experiments to gain more detailed insight into the folding free-energy landscape of ADE and its unfolding/refolding pathway. In constant velocity experiments the prominent intermediate described above is clearly visible (**Fig. S1A**, I1, green WLC fit). However, a zoom-in of the trace reveals additional intermediates with contour lengths both above and below I1 (**Fig. S1A**). A passive mode measurement (see Material and Methods) allows the analysis of the exchange kinetics between the various states via folding/unfolding fluctuations over time (**Fig. 2B**). We observed multiple close-to-equilibrium transitions between various states over many seconds. A zoom into this trace (**Fig. 2B**) allowed for observing the folding/unfolding pathway in real time. According to their different contour lengths, we identified six different levels. Based on those lengths, we ran a Hidden Markov Model (see SI and(27) for more details) (HMM), which yielded an assignment of the data points to the various states (**Fig. 2B**) as well as the states’ lifetime distributions (**Fig. S4A**). In sum, the ADE fluctuates between the fully folded state (F) and the completely unfolded state (U) (**Fig. 2 B and C**). While most states can directly pass to their neighboring ones, the orange state can only be populated from and depopulate into the red state, but we never observe a direct transition from the blue or green states into this state, or vice versa. This indicates that the orange state is an off-pathway intermediate, unable to productively fold further.

Hence, we identified the orange state (M) as a misfolded state, while all the other intermediates are on-pathway: I1 (green), I2 (red) and I3 (purple). From the change in force upon each transition, we calculated the associated gain in contour length (**Table S1**) and suggest structural models for the respective intermediates (**Fig. 2C**). To define the structural model of the on-pathway intermediates, we assumed that each individual unfolding step opens a stem region together with the subsequent interior bulge. The structures shown in **Fig. 2C** satisfy all constraints. For the structure of the misfolded intermediate (M), we used the information of its contour length as well as its exclusive accessibility from I2. Free energy differences at zero force between all the states assigned in passive mode are summarized in **Table S2** (see SI for more details).

Analysis of the lifetimes of the different states hints towards more transient states constituting the folding pathway. While the lifetime distributions of the F, U and I1 are single exponential, the distributions of I2, I3 as well as M are multi-exponential (**Fig. S4A**). This finding suggests that those states are, in fact, a mixture of different states occurring at similar contour lengths.

We can, therefore, extend our model for the folding/unfolding pathway of ADE as shown schematically in **Fig. 2D**. While the on-pathway intermediates directly connect the native to the unfolded state, there are several branch points where misfolds can slow down the folding process. The misfold M could be directly identified owing to its distinct length, while the other misfolded intermediates have a length indistinguishable from the Intermediates I2 and I3. We hence denote them as m_I2_ and m_I3_. Detailed suggestions for structural models of the misfolded intermediates are given in **Fig. S4B**. As a summary, the ADE samples a large conformational space, while its folding equilibrium is shifted towards the full stem-looped structure.

### Roquin stabilizes the apical stem-loop structure of the *Ox40* ADE

To assess the impact of Roquin onto the ADE stability, we next studied binding and unbinding of Roquin to the *Ox40* ADE in single-molecule experiments under near-physiological buffer conditions (low salt). For comparison and alignment to the structural features determined in high salt buffer, a microfluidic chamber allowed us to expose the same molecule of the *Ox40* ADE to up to three different buffer conditions (**Fig. S5**) as shown in **Fig. 3A**, displaying stretch/relax cycles in high and low salt buffer. Even though forces and stability in low salt are lower compared to high salt, we observe a conservation of the essential features of the unfolding pathway. Note that the decreased stability at low salt precluded clear identification of state I2 owed to the inherent loss of resolution of optical traps at lower forces, and I2 is therefore not annotated anymore. Binding of coreROQ manifests itself in a distinct stabilization of I3, in line with previously observed stabilization of CDEs(2), while adding extROQ leads to an even higher stabilization of I3. Importantly, both coreROQ and extROQ exclusively stabilize I3, representing the apical stem-loop structure of the *Ox40* ADE (basepairs made up of residues 70-73 and 80-83, **Fig. 1A**), but no other parts.

**Figure 3.**
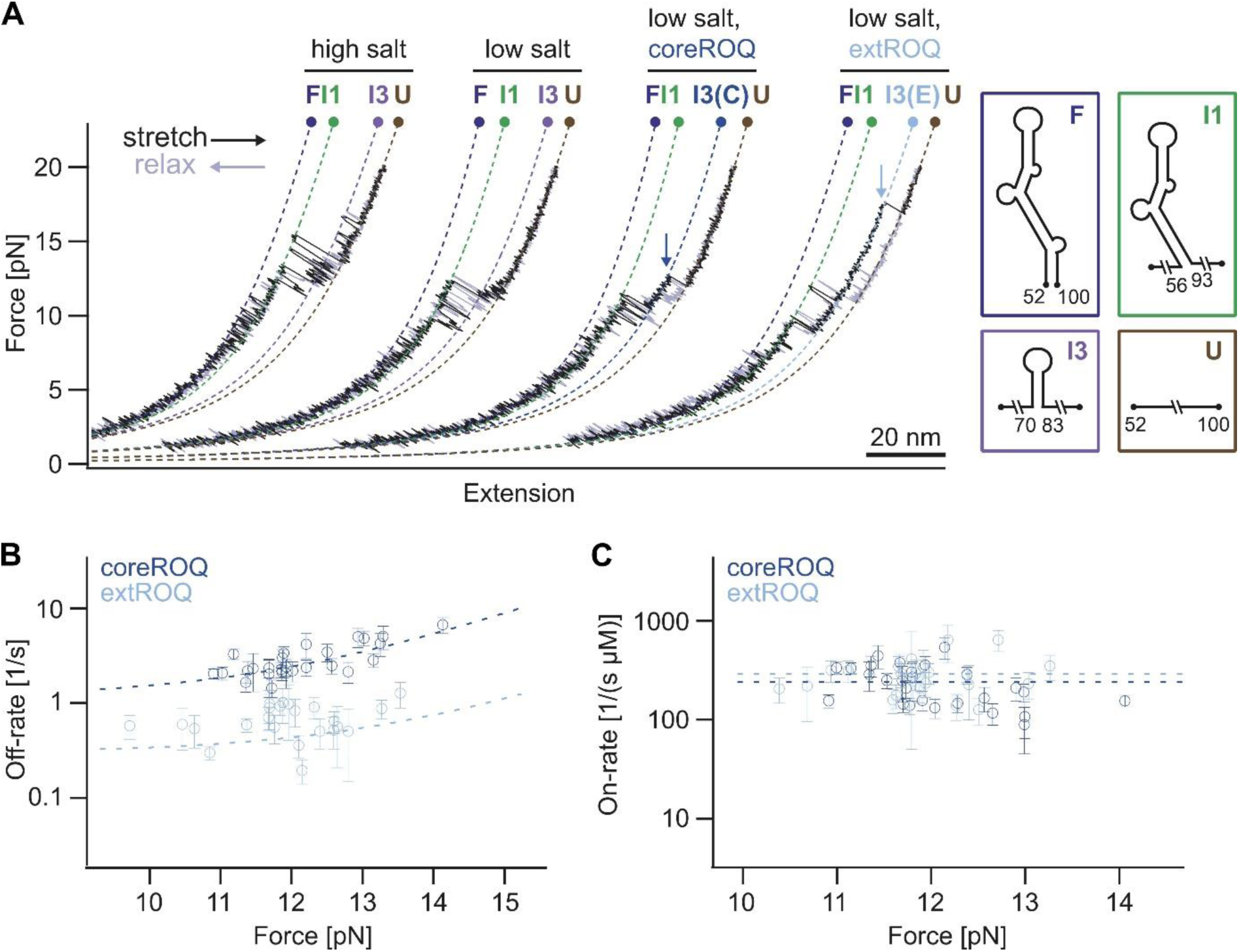
Roquin stabilizes the apical stem-loop of the *Ox40* ADE. **A)** Representative force-extension curves of the ADE in different conditions: high salt (HS), low salt (LS), LS + coreROQ, LS + extROQ. Note that the extension axis does not display absolute values since the traces are arranged for side-by-side comparison. The measurements of the ADE with coreROQ and extROQ reveal stabilization of the same I3 intermediate (unbinding and unfolding events marked by dark blue (coreROQ) and light blue (extROQ) arrows) by both proteins. I3 (C) = coreROQ bound I3, I3 (E) = extROQ bound I3. **B)** Unbinding rates of coreROQ (N = 27) and extROQ (N = 23) from ADE in LS buffer, extracted from passive-mode experiments and plotted against the force of the bound state in the respective traces. An exponential fit (dashed line, see **Table S3** for fitting parameters) was applied, assuming a force-dependent and a force-independent contribution to the off-rate. **C)** Binding rates of coreROQ (N =27) and extROQ (N = 23) to a folded ADE structure (see Materials and Methods for details), extracted from passive-mode experiments. A force-independent constant fit (see **Table S3** for fitting parameters, dashed line) was applied to the data points.

Analysis of on- and off-rates in passive-mode experiments (**Fig. S6, Fig. S7**) showed that the increased stabilization through extROQ binding compared to coreROQ is driven by reduced off-rates, while on rates are identical (**Fig. 3B** and **C**). The measured on-rates are not force-dependent (**Fig. 3C**), suggesting that the affinities of both coreROQ and extROQ to folded I3 are significantly higher than to unfolded RNA (see Materials and Methods for more details). Based on the measured on- and off-rates (**Table S3**), we calculated an affinity at zero force of 4.8 ± 0.4 nM (coreROQ) and 1.03 ± 0.14 nM (extROQ). In summary, these data provide evidence for an RNA hairpin-stabilizing effect of Roquin, enhanced by lowered off-rates mediated by the Roquin B-site.

### Roquin promotes destabilization of lower stem regions

As the ADE forms a structurally closed entity, the effect of Roquin onto the apical stem-loop and the basal structured segments needs to be assessed in the full ADE context. We first recorded CD melting curves of the ADE alone and in complex with core or extROQ (**Fig. 4A**). The apo ADE showed a melting point at 76.7 °C (T_m2_), in line with(7). Besides we observed a less pronounced melting event at around ∼65 °C (T_m1_), which could not be fitted. This initial transition, which was not observed before, is likely due to a local modulation of stability in the physiological buffer compared to (7). To our surprise, in presence of the proteins we observed a decrease for both melting points. Strikingly, T_m1_ showed a stronger reduction than T_m2_, and the effect was more pronounced during binding of extROQ (**Fig. 4A**). Roquin hence exhibits a destabilizing effect on parts of the RNA duplex, and the B-site actively contributes to this. This is also reflected in an overall decreased A-type helical content, observable by a lowered absolute starting signal of ellipticity in CD(28) (**Fig. 4A**, **Fig. S8A**). Interestingly, the complex melting points are higher than the apo Roquin melting points, suggesting a stabilizing effect of the RNA onto Roquin (**Fig. S8B**).

**Figure 4.**
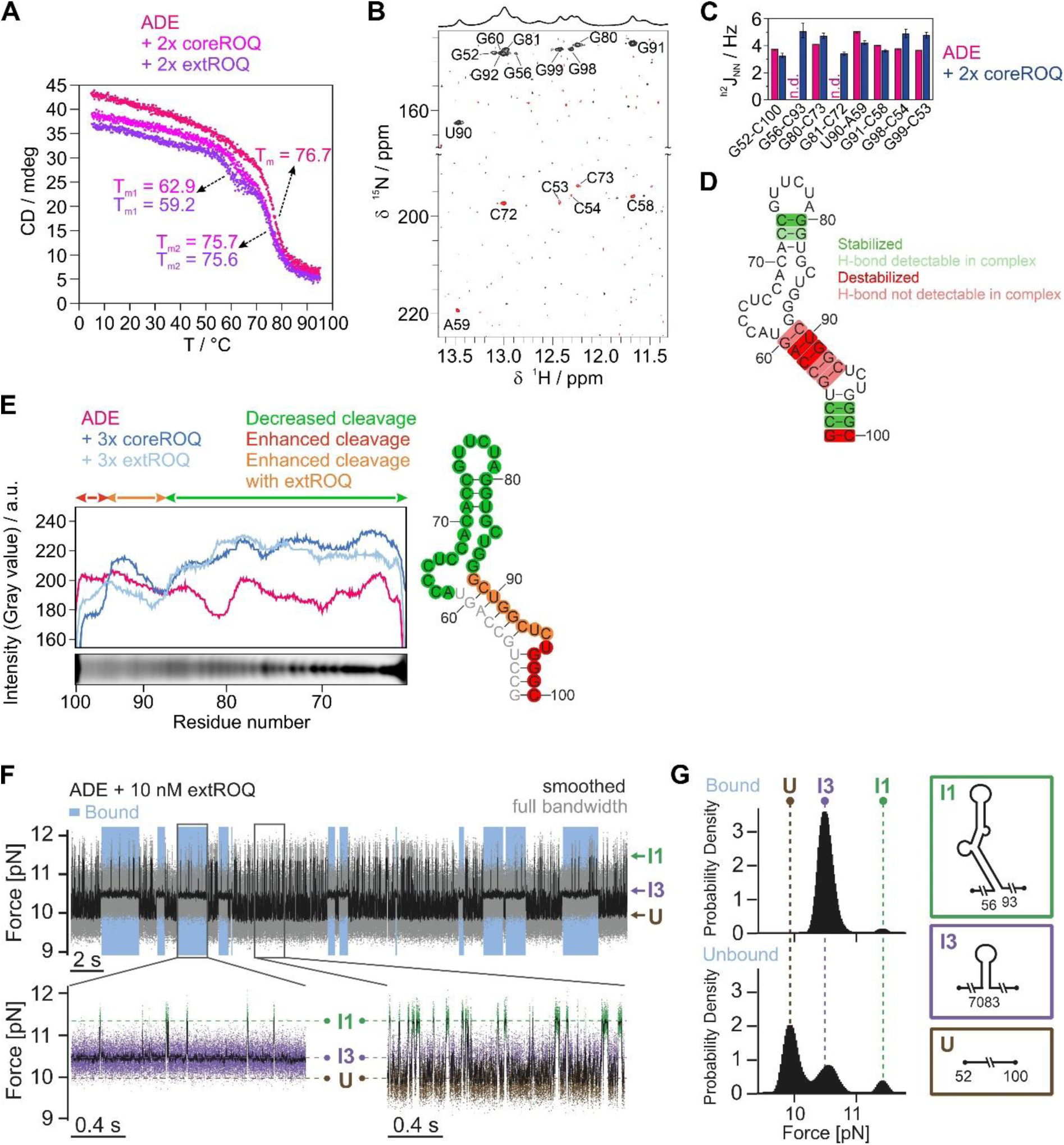
Roquin mediates destabilization of the lower *Ox40* ADE stem. **A)** CD melting curve of apo ADE and ADE bound to coreROQ and extROQ. Individual melting points are indicated. **B)** HNN-COSY spectrum of the ADE in complex with coreROQ. H-bond donor peaks are shown in black, pairing H-bond acceptor peaks in red. The corresponding 1D spectrum is shown above. **C)** Quantification of H-bonds observed in HNN-COSY experiments of apo and complexed ADE shown in Fig. 2A and 4B. **D)** Base pairs stabilized or destabilized by Roquin are highlighted in green and red, respectively. Faint green and red indicate bases that cannot be quantified in either of the two spectra. **E)** Hydroxyl radical probing of apo ADE and ADE bound to coreROQ and extROQ. Cleavage profiles corresponding to the PAGE shown in **Fig. S8C** are shown aligned to the alkaline ladder below. Colored arrows indicate regions with decreased or increased cleavage of the protein bound ADE compared to the apo RNA. Differences in cleavage are plotted onto the ADE secondary structure on the right. **F)** Top, passive-mode trace of the ADE with 10 nM extROQ at a lower force regime where ADE populates folded structures up to I1. The protein bound states (highlighted in light blue) can be discerned by strong stabilization of I3 and lower population of more folded structures. Bottom, representative zoom-ins to the bound (left) and unbound (right) phases. **G)** Direct comparison of population probabilities in the extROQ bound (upper) and unbound (lower) case. The histograms display the distribution of the force data and show the different population probability of states U, I3 and I1 (indicated in dashed lines, corresponding structures on the right), collected from the bound and unbound phases of the passive-mode trace shown in F).

The CD melting data can only yield global readouts of regional stability, not considering nucleotide-resolved differences. Still, they suggest the existence of RNA regions with different responsiveness to binding by Roquin. To track effects on the level of single basepairs, we recorded an HNN COSY NMR(26) spectrum of the coreROQ-bound ADE (**Fig. 4B**). While H-bonds were still found, the reduction in signal intensity especially for H-bond acceptors indicates an overall destabilizing effect of Roquin on the monitored duplex regions. We quantified the H-bonds of the apo and complex state (**Fig. 4C**) and observed an increase in H-bond strength for base pairs in the apical stem loop and the basal stem (**Fig. 4D**), pointing at elevated stability. Interestingly, base pairs with reduced H-bond strength, and thus stability in presence of Roquin cluster in the stem region below the Bulge (bp 56-93 until bp 60-89, **Fig. 4D**). To corroborate our findings and to obtain information of non-base paired residues, we performed hydroxyl radical probing (OH• probing) on the ADE in the apo and Roquin-bound form. As OH• probing reports on solvent accessibility of RNA, it can be used to track conformational changes within footprinting experiments(29). Cleavage profiles extracted from denaturing PAGE (**Fig. S8C**) showed decreased cleavage of residues 62-87 in presence of either core or extROQ compared to the apo ADE (**Fig. 4E**). For residues 88-100 we observed increased cleavage, while the PAGE-limited resolution did not allow quantification of bands corresponding to residues 52-61. Upon mapping the changes in cleavage onto the secondary structure, we concluded that Roquin not only stabilizes the apical stem-loop, but at least protects the adjacent stem structure and the Bulge from cleavage. In contrast, the entire structure below the Bulge is more solvent exposed and destabilized. Our NMR data and OH•probing experiments are thus in good agreement with CD melting curves and suggest that T_m1_ corresponds to melting of the lower ADE half. The upper half is stabilized through Roquin binding, likely accompanied by modulation of the RNA fold. This rationalizes the slight yet detectable decrease of T_m2_, while we locally observe stronger H-bonds in NMR. Hence, we claim that Roquin exhibits opposite effects on the stability of distinct stem regions.

### The Roquin B-site shifts RNA structural equilibria to partially opened conformations

To investigate the influence of Roquin on the stability and folding kinetics of the individual ADE substructures, we next recorded passive-mode folding traces in the presence of extROQ at lower forces (**Fig. 4F**, **Fig. S9** and **Fig. S10**). In the full-view (**Fig. 4F**), bound and unbound states can be clearly distinguished: while binding of extROQ has a pronounced stabilizing effect on I3 and prevents population of the unfolded state, the unbound states show rapid fluctuations between I3 and U. In addition to the expected stabilization through extROQ binding, we observe that during bound phases, further folded states are less populated as compared to unbound phases (i.e. less transitions from I3 to I1). As noted above, I2 and I3 cannot be reliably distinguished from each other at the low salt conditions and both populations are lumped into I3. Zoom-ins to bound and unbound phases (**Fig. 4F**, bottom) clearly showed this discrimination of the more folded structure (I1) relative to the less folded structure (I3) by extROQ. **Fig. 4G** directly compares the population distributions. The differences in population yielded a destabilization of I1 by 1.3 ± 0.4 kcal mol^-1^ by interaction with extROQ. Analysis of folding/unfolding rates between I1 and I3 in presence and absence of extROQ (**Fig. S10F** and **Table S4**) showed that extROQ predominantly slows the refolding rate while leaving the unfolding rate unaffected. Analysis of the population shift upon addition of coreROQ also showed a destabilizing effect on I1, albeit much smaller (0.34 ± 0.05 kcal mol^-1^, **Fig. S10D**). Similar to extROQ, the effect of destabilization through coreROQ also acts on the folding rates rather than unfolding rates (**Fig. S10E** and **Table S4**). Altogether, we propose a twofold effect of Roquin on the ADE structure: stabilization of the apical stem loop by binding of the A-site and destabilization of the basal stem below the Bulge by interaction of the B-site.

### The Roquin B-site binds ssRNAs

A simple mechanism explaining the destabilization of the secondary structure between I3 and I1 through slowing of the refolding rate would entail interaction of parts of coreROQ and extROQ with single-stranded parts of the involved ADE region. We thus tested both proteins for interactions with the single-stranded halves of the ADE. We also included an RNA duplex annealed from the two halves, in order to probe for potential double-stranded (ds) RNA interactions as suggested by Tan et al.(6). Imino proton NMR spectra confirmed the single-stranded nature of the ADE halves as well as duplex formation for the annealed species (**Fig. S11**). In electrophoretic mobility shift assays (EMSAs) we did not observe binding of coreROQ to either of these RNAs (**Fig. 5A**), while extROQ interacted at low-micromolar concentrations with both single-stranded ADE halves and the duplex. This was confirmed by ^1^H,^15^N-HSQC NMR spectra (**Fig. 5B**): In presence of ss or dsRNA only minor chemical shift changes were observed for coreROQ (**Fig. 5B**, upper row). ExtROQ, however, exhibited larger chemical shift perturbations (CSPs) and line-broadening for the single-stranded ADE halves and the duplex RNA (**Fig. 5B**, lower row), confirming our EMSA results. Interestingly, in extROQ NMR experiments also signals assigned to the coreROQ domain were affected by RNA binding, suggesting that both A- and B-site contribute to recognition of ss and dsRNA, but that the B-site is the main driver of interaction. Roquin is thus able to not only interact with hairpin structures, but also recognizes ss and dsRNAs through its B-site.

**Figure 5.**
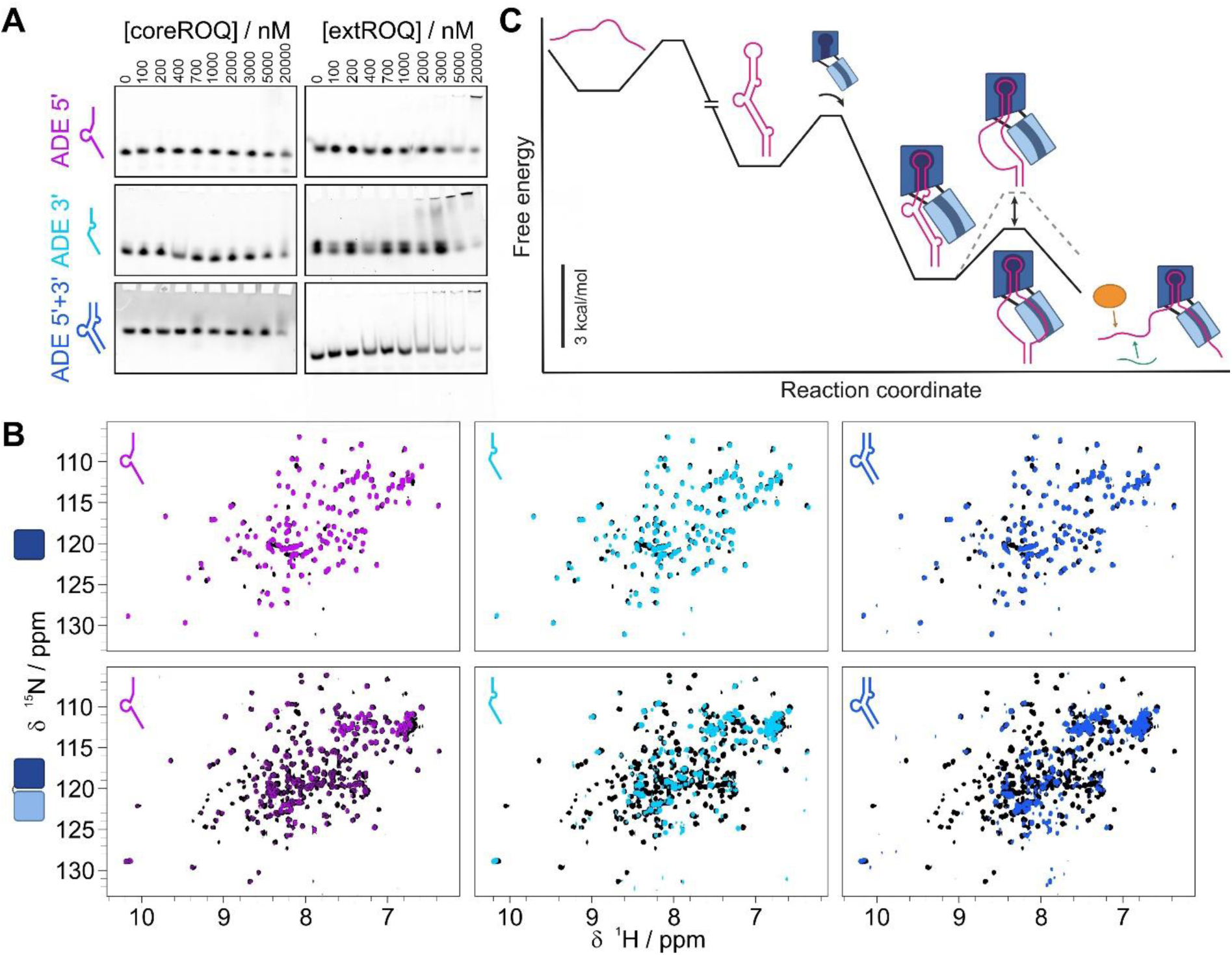
Roquin interacts with ss and ds RNAs through the B-site. **A)** EMSAs of coreROQ and extROQ with ss ADE 5’ and 3’ halves and an annealed duplex. Protein concentrations are plotted on top. **B)** ^1^H,^15^N-HSQC spectra of apo (black) coreROQ (top) and extROQ (bottom) and in complex with the ss ADE 5’ half (purple), ss ADE 3’ half (cyan) and the ds ADE 5’+3’ of the central stem (blue). **C)** Model of the energy landscape for extROQ binding the ADE. In a first step, ADE folds into its native structure. Next, extROQ engages with the apical stem loop through its A-site. Binding of the B-site to ssRNA reduces the barrier for partial unwinding of the central stem regions by shifting the equilibrium towards a partially opened ADE. Destabilization of ADE stem structures facilitates *trans* factor engagement or enhances Roquin binding by conformational adjustments. The relative differences between energy minima are quantitative (assuming a 1 µM concentration of extROQ) while barrier heights are not.

We wondered whether sequence composition or geometry of the ADE stem affects RNP formation. To test this, we created an artificial ADE where we mirrored the stem, but maintained the loop orientation (**Fig. S12A**, called ADE_mir_). Two mutations within the Bulge were required to stabilize the properly mirrored ADE geometry, and structural integrity was confirmed by a ^1^H,^1^H-NOESY NMR spectrum (**Fig. S12A**). In EMSAs both coreROQ and extROQ showed reduced binding to the ADE_mirr_ (470 and 458 nM, respectively) compared to the wt ADE (307 and 322 nM, respectively) (**Fig. S12B** and **C**). Smeary complex bands further suggested that the complex was less stable in a gel matrix. NMR ^1^H,^15^N-HSQC spectra of coreROQ with wt ADE and ADE_mirr_ showed characteristic CSPs (**Fig. S12D**) confirming complex formation, while extROQ complex spectra suffered from severe line broadening due to increased molecular weight and chemical exchange. Both coreROQ complex spectra were virtually identical (**Fig. S12E**), suggesting a conserved ADE-like binding mode for the mirrored ADE version. We thus conclude that the mirrored ADE provides a less favorable geometry for RNP formation, resulting in lowered on-rates, but that Roquin induces an ADE-like fold for engagement, as suggested before(8).

## DISCUSSION

The spatiotemporal availability of an RNA *cis* element’s functional conformation determines its engagement with cognate *trans*-acting binding partners, e.g. proteins. Despite its limited chemical space, RNA exhibits a great structural heterogeneity and plasticity, also within RNPs. Small iterations in RNA sequence, and hence structure, can alter the RNA interactome landscape and lead to misfunctional (and pathogenic) RNPs. Consequently, we strive for a detailed understanding of RNP structures and mechanisms of complex formation as a basis for gene regulation or as the causative of diseases. However, detailed structural and mechanistic insights of plastic RNPs are challenging to obtain holistically, and available methods often fail to capture more than individual aspects in the full description of an RNP.

### The combination of OT and NMR captures and quantifies RNA/RNP folding landscapes

We here combined NMR spectroscopy with single-molecule OT experiments and RNA-protein biochemistry to describe I) the folding landscape of a medium-sized RNA element, II) the specific engagement of two interrelated, non-redundant RBDs of the protein Roquin with distinct RNA element regions, and III) how Roquin exploits its multi-domain architecture to modulate the RNA folding landscape. We integrated the high-resolution structural data from NMR with time-resolved single-molecule information from OT. We show that OT experiments provide single-nt information on a sub-ms timescale for RNAs varying from 29 nt to 140 nt (**Fig. 1-4**). NMR can help to assign folding events in OT traces. On the other hand, the low amount of material required for OT experiments allows to test RNA folding under varying conditions to select promising features for *en detail* testing in NMR. The low concentration also allows suppressing unspecific intermolecular influences, often observed for RNAs, e.g. based on stacking interactions(30, 31) or stem-loop intrinsic complementarity, even at small size(32) .

Due to its inherent structural flexibility(7) the *Ox40* 3’UTR shows overlapping unfolding profiles, which we could reconstitute in a divide-and-conquer approach (**Fig. 1C**). OT experiments allowed to distinguish cooperative one-step folding/unfolding (CDE) from more complex multistep folding with multiple intermediate states found in ADE, ADE-CDE and the fl construct. While NMR experiments confirmed a network of hydrogen bonds as the basis for a major stable conformation of the ADE (**Fig. 2A**), OT revealed a complex dynamic folding network with stable on and off-pathway intermediates (**Fig. 2B-D**). Bulk measurements usually capture a population- and time-averaged conformation, whereas single-molecule experiments allow to dissect low-populated folding states more directly and with a much higher sensitivity. This now allowed to monitor unfolding of the ADE nts 67-69 (I2 in **Fig. 2B, C**), a region previously invisible in NMR experiments(7), as well as distinguishing between two alternative structure predictions of the fl *Ox40* 3’UTR (**Fig. 1C**).

Multiple RNA folding intermediates and their constant exchange can provide target sites for multiple RBPs with varying specificities as shown before(2). The multiple intermediates we find for the *Ox40* ADE show the great plasticity RNAs can exhibit. How a single sequence can translate into heterogeneous structures could possibly explain inconsistencies between *in vitro* and *in vivo* data for RNA folds, as e.g. found by Andrzejewska et al. (33), and their susceptibility to fold-mediated interactions with RNA-binding proteins.

A valuable byproduct of OT measurements is quantitative information about thermodynamics and kinetics. In combination with microfluidics, we were able to provide single-molecule affinities and on/off-rates for complex formation between ROQ and ADE. Our observed affinity of extROQ was significantly higher (∼170-fold) compared to previously reported bulk measurements(7), obtained from EMSA (171 nM). Although useful for qualitative comparison, EMSAs often do not yield absolute affinities since they are carried out under non-equilibrium conditions. Here, single-molecule measurements offer an attractive alternative. In this study, the resultant availability of kinetic data could readily explain the enhanced stability of extROQ-ADE compared to coreROQ-ADE as off-rate driven.

We further showed that OT data can fully complement the gaps in ADE RNA coverage, found as technical limitations in NMR and biochemical probing (**Fig. 4A-E**). For the Roquin-ADE complex, OT experiments not only agree well with NMR, CD and structure probing data, but also allowed quantification of RNA conformational populations. We suggest this methodology is easily but efficiently adaptable to other highly specific RNA-protein complexes. In similar studies, an integrative approach using NMR and single molecule FRET experiments had already been successfully taken to measure protein/protein and protein/RNA binding kinetics(34, 35). Jones et al.(36) have used NMR combined with OT to study microRNA structure and dynamics. While OT experiments in that study were limited to (un)folding of RNA, our combination of microfluidics-enhanced OT with NMR now reveals the energy landscape of protein/RNA interactions. It provides unique access to hidden information in RNA folding pathways obtainable from low sample amounts and possible to visualize at atomic level.

### Roquin modulates RNA structures in a bimodal fashion through A and B-sites

The experiments in **Fig. 4** suggest that, apart from the apical stem-loop, Roquin also interacts with other parts of the ADE. While this effect is small for coreROQ, binding of extROQ clearly destabilizes the ADE structure basal to the apical stem-loop. Our combined approach exclusively allows us to pinpoint the affected region to the central stem between residues 56-60. How can we explain the B-site-induced destabilization of the central stem? In summary, our data support the sequence of events outlined in **Fig. 5C**. After folding, extROQ binds strongly to the apical stem loop through its A-site. Even though we found that the Roquin B-site can bind both ss and dsRNA with a similar, low µM-affinity (**Fig. 5A**) the observed reduction in central stem stability suggests that extROQ favors binding to ssRNA for efficient engagement with the full ADE. This preference could be supported by structural constraints occurring with the A-site tightly bound to the apical stem loop and the B-site trying to engage with the adjacent central stem. In this scenario, unfolded flexible ssRNA strands, rather than duplexed RNA, may facilitate proper engagement. Note that the conformation of an A-site-only engagement is thermodynamically favored over the fully engaged conformation (**Fig. 5C**) because the energetic costs of opening the central stem (see **Table S2**) outweigh the free energy gained by B-site binding (see **Fig 4G**). However, B-site engagement to ssRNA facilitates the opening of the central stem necessary for the binding of downstream effectors. It both reduces the kinetic barrier for the separation of the two RNA strands and shifts the conformational equilibrium of the central stem towards unfolded (**Fig. 5C**). In this context, the interior bulge involving nt 62-66 may act as a seed to initiate ssRNA binding, thus facilitating splitting of the double strand. The exact mechanism remains to be explored, but our EMSAs suggest a moderate preference for the ADE 3’ strand engaged by the B-site.

The new role for the B-site in Roquin as a ssRNA binder expands its target spectrum beyond hairpins and dsRNA. In line with previous findings, the ssRNA binding capacity of Roquin might have served in target selection during evolution(8) and be of particular relevance in unstable element binding. In this, the ROQ domain B-site could have enhanced affinity to otherwise less favorable targets, i.e. with unstable hairpins or stem-loops of suboptimal geometry. Thereby the rather promiscuous binding of the B-site potentially compensated for weaker A-site binding.

As ADEs differ significantly from CDEs in structure(8) and show a distinct folding behavior (this study) we speculate that structural features like the central ADE Bulge are required for modulation by Roquin also in other targets, i.e. to initiate unwinding. Possibly, Roquin modulates RNA *cis* element stability to reach an optimal window for subsequent engagement. At the same time, this mechanism exposes the newly ss, i.e. now unpaired region of the stem for possible interactions with *trans* factors, be it microRNAs or proteins, that could contribute to regulation by Roquin (**Fig. 5C**), e.g. in acting as target sites for recruited nucleases.

With respect to posttranscriptional regulation by Roquin, we suggest that specific target mRNA recognition and suppression additionally involve RNA sequence-encoded dynamics and shift-in-equilibrium caused by the protein. Those remain challenging to put into numerical descriptors. But it appears obvious that a canonical target sequence of Roquin, besides the previously described stem-loops caught by coreROQ (including recent specification of loop requirements(37)), needs to take into account the differential affinity of the extROQ B-site to an adjacent RNA region as well as its intrinsic plasticity and response to ROQ-binding. Further, a full understanding of Roquin-mediated gene regulation will require to understand the effect of Roquin interactions with other proteins(38, 39), and potential Roquin cooperativity or homodimerization exploiting the occurrence of multiple *cis* elements in a 3’UTR(40).

In summary, our approach allowed precise detection and quantification of folding intermediates in a dynamic RNA, both alone and in complex with a two-domain RBP. We provided mechanistic insights in how two RBDs exhibit opposing effects on RNA structure stability and plasticity to fine-tune complex formation, representing a specificity-determining parameter.

## MATERIAL AND METHODS

### Protein expression and purification

The ROQ domain of murine Roquin-1 was expressed as two constructs as described before(7): coreROQ and extROQ were expressed in *E. coli* BL21 at 37 °C overnight while shaking. For isotope labelling minimal M9 medium was used supplemented with ^15^NH_4_Cl. Briefly, cells were harvested, sonicated and proteins were purified via IMAC (immobilized metal ion affinity chromatography). After TEV protease (tobacco etch virus) cleavage overnight, proteins were passed through a reverse IMAC. The flow-through was concentrated and purified via SEC (Superdex S75 16/600 from GE). Proteins were buffer exchanged to 150 mM NaCl, 50 mM Tris pH 7.0 and 1 mM TCEP. Purity and structural integrity of the proteins were checked by SDS-PAGE and NMR spectroscopy.

### RNA *in vitro* transcription and purification

Templates for *in vitro* transcription of RNAs for OT experiments and mirrored ADE were generated by PCR and Gibson assembly of target sequences into HDV-containing vector (Hepatitis D virus ribozyme). Wt CDE, ADE tandem ADE-CDE and fl *Ox40* 3’UTR constructs were already available from(7). RNAs were transcribed and purified as before(7), i.e. after transcription RNAs were precipitated, purified via a denaturing urea polyacrylamide gel, eluted by ‘crush-and-soak’, buffer exchanged and stored at – 80 °C until used. RNAs used for OT experiments (*Ox40* full-length (fl), tandem ADE-CDE (ADE-CDE), ADE) were transcribed with 5’ and 3’ overhangs to anneal with DNA handles (**Table S5**). Note that the OT fl construct contained residues 1-140, omitting the flexible 3’ tail. For NMR, CD and EMSA experiments RNAs were snap-cooled prior use. Single-stranded (ss) 5’ and 3’ fragments of the ADE were obtained from Horizon and dissolved in the protein buffer (see above). The annealed duplex (ds) was formed by mixing the 5’ and 3’ halves in equimolar amounts, incubation at 95°C for 5 minutes and a slow cool-down to room temperature on the bench.

### NMR spectroscopy

All NMR experiments were performed on Bruker Avance III Spectrometer operating at 700 and 950 MHz proton lamor frequency equipped with triple-resonance cryogenic probeheads. For measurements and processing of spectra Topspin versions 3 and 4 were used. Spectra were analyzed with NMRFAM-Sparky 1.414(41).

HNN-COSY(26) experiments were recorded as a BEST-TROSY version on a 275 µM ADE sample for the apo state and in presence of a two-fold molar excess of coreROQ at 283 K. The transfer time was 30 msec, and 384 indirect points were collected for the apo RNA and 256 for the complex. H-bond J-couplings were calculated according to

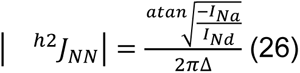

where I denotes the intensity of the H-bond donor (Nd) or acceptor (Na) and 2Δ is the COSY transfer time.

^1^H,^15^N-HSQC spectra were recorded on uniformly labeled ^15^N apo coreROQ and extROQ proteins and in presence of a 1.2-fold molar excess of wt or mirrored ADE and a 3-fold molar excess of ss or dsRNAs. A ^1^H,^1^H-NOESY of the mirrored ADE was recorded at 283 K on a 300 µM sample with 416 indirect points and a mixing time of 250 msec.

### OT measurements

A commercial microfluidic chip (C-Trap® Optical Tweezers - Fluorescence & Label-free Microscopy (LUMICKS)) comprising two bead channels and three flow channels that can maintain separate phases through laminar flow was used, allowing the measurement of the same molecule under different conditions (**Fig. S5**). To ensure correct attachment of sample constructs to one streptavidin (SV) and one anti-digoxigenin (AD) coated silica bead, SV and AD beads were inserted into separate channels, allowing selective trapping of the two bead types by each of the respective trapping lasers. Assembled constructs were incubated at room temperature for 10 min with SV beads beforehand. This solution and the AD beads were then both diluted to a bead density appropriate for optical trapping of individual beads in 300 µl of HS buffer. For measurements with protein, two of the flow channels were run with HS and LS buffer to ensure correct tethering, and LS buffer with varying concentrations of coreROQ or extROQ was run in the third channel. Oxidative damage was counteracted by addition of a GLOXY scavenger system (see SI for more details) to all five bead and measurement solutions, which were then injected into the optical trap setup (C-Trap® Optical Tweezers – Fluorescence & Label-free Microscopy) via syringe pumps.

The dual-beam OT setup was assembled by successively trapping an SV bead, pre-incubated with the sample construct, and an AD bead into a respective laser and bringing them into close proximity, allowing establishment of a construct tether. Initial quality control via fingerprint stretch-relax cycles in HS buffer was followed by measurement in the same or a different channel, according to the specific interest of the experiment. To extract quantitative kinetic and energetic information, passive mode experiments were performed, in which the distance between the two laser beams was held constant over time. This allowed the collection of equilibrium unfolding/refolding and binding/unbinding data. Measurements were performed at ≈ 25 °C.

### Rate determination

To extract on- and off-rates of coreROQ or extROQ to the ADE, passive-mode traces were HMM assigned(27). As ROQ binding drastically increases the lifetime of I3 and higher folds, bound and unbound states can easily be discriminated. The off-rate *K_off_* corresponds to the inverse of the average lifetime 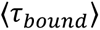 of ROQ-bound states:

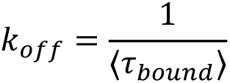

Since the off-rates display a force dependence, an additional force-induced unbinding mechanism can be presumed. The off-rate can then be modeled by a force dependent (first summand) and a force independent (second summand) contribution:

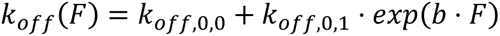

where *k_off,0,0_* designates the force-independent contribution to the off-rate, while parameters *k_off,0,1_* and *b* define how strong the force influences the off-rate. The off-rate at zero force correspondingly calculates as 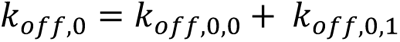

As the on-rate *k_on_* expresses binding to an apical stem-loop ADE structure, i.e. to I3 or more folded intermediates, its calculation has to take into account the force-dependence of the folding population distribution of ADE. The effective on-rate decreases with increasing force, because the protein-accessible not-unfolded states are defavorized against the unfolded state. To calculate the on-rate 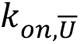 of coreROQ or extROQ to the not-unfolded state of ADE (from I3 onwards), the inverse of the average lifetime of unbound states 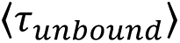 at a given force bias has to be factored with the relative probability of the not-unfolded 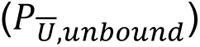 to the total 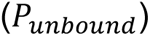 population of this state. Together with division by the protein concentration [*R*], this yields:

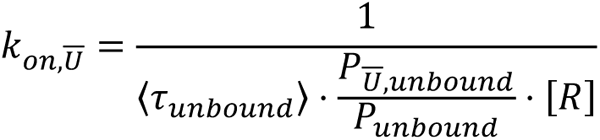

This rate does not show a significant force dependence and the on-rate at zero-load 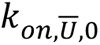 can thus be calculated as the average of 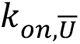.

### Circular dichroism spectroscopy

CD spectra were recorded on a JASCO J-810 spectrometer in the protein buffer from 180-320 nm. 20 µM ADE RNA were measured in presence of 0, 5, 10, 20 and 40 µM of coreROQ or extROQ. For melting curves, the changes in the CD signal at 263 nm was monitored from 5 – 95 °C with a temperature gradient of 1°C/min for 20 µM apo ADE RNA or in presence of 40 µM of the respective protein. Curve fitting for separated T_m_s was done in Origin with DoubleBoltzmann fits.

### Electrophoretic Mobility Shift Assay

For EMSAs RNAs were fluorescently 5’-labeled with fluorescein as described before(7). EMSAs were performed according to (42). Briefly, 3 nM of ADE RNA were incubated with 0, 100, 200, 400, 700, 1000, 2000, 3000, 5000 and 20000 nM of coreROQ or extROQ. Complexes were run on a native 6% polyacrylamide gel at 4°C for 1 hour. Gels were imaged with a ChemiDoc Imager (BioRad). Quantification of gel bands was performed with ImageJ and K_D_s were calculated as averages plus standard deviations from triplicates fit individually with a Hill1 fit in Origin.

### Hydroxyl radical probing of RNA

60 pmol of fluorescently labeled ADE RNA apo or in presence of a 3-fold molar excess of coreROQ or extROQ were rapidly mixed with 1 mM Fe(II), 2 mM EDTA, 1 mM sodium ascorbate and 0.06% (w/w) H_2_O_2_ according to (43). After 3 minutes of incubation, samples were quenched with 10 mM thiorurea and loaded onto a denaturing 10% polyacrylamide gel (7). For imaging a Typhoon 9400 Variable Mode Imager (GE) was used and gray scale intensity profiles for all lanes were extracted with ImageJ. These profiles were plotted against the nucleotide residue number obtained from the alkaline ladder. Alkaline and T1 ladders were generated as described before(7).

## ACCESSION NUMBERS

## ACKNOWLEDGEMENT

We acknowledge lab members of the Schlundt and Rief labs for critical discussions. We thank Katharina Targaczewski, Ulrike Majdic and Lion Reichardt for technical support and Anna Wacker for providing the HNN-COSY pulse program. The Frankfurt BMRZ acknowledges support from the state of Hesse and funding via the IWB-EFRE-program 20007375.

