## Supplementary Information for "Roquin exhibits opposing effects on RNA stem-loop stability through its two ROQ domain binding sites"

### EXPERIMENTAL PROCEDURES AND DATA ANALYSIS

The experimental procedures used for optical tweezer experiments and the analysis methods of the data are described in the following.

#### DNA handle preparation and hybridization reaction

Synthesis and purification of DNA handles has previously been described in Walbrun et al.(1). Biotin and digoxigenin-functionalized handles were PCR-amplified separately using a lambda-phage template DNA, forward primers with biotin or digoxigenin modifications, and backward primers introducing ssDNA overhangs (all sequences are listed below in **Table S5**). Primers were supplied by Biomers. Water was ultrafiltrated by Sartorius arium pro UF. Other components for the PCR reaction were ordered from NEB. 300 µl of reaction mix (1x Thermopol buffer, 0.2 mM dNTPs, 6 µl lambda-phage DNA, 1 µM forward (biotin/digoxigenin) and backward primers, 1 µl Taq polymerase) were set up and split equally (50 µl) into six PCR reaction tubes, run in the peqSTAR thermocycler from PEQLAB (for thermal protocol see **Table S6**). PCR products were purified using the Monarch PCR & DNA Cleanup Kit (NEB), using the hybridization buffer (300 mM NaCl, 50 mM Tris (pH = 9.0), 20 mM EDTA (pH = 8.0)) for elution. Quality of isolated products was verified *via* agarose gel electrophoresis (1.5%) using TAE buffer (40mM Tris, 20mM Acetic acid, 1mM EDTA).

Assembly of the entire construct used for measurement, comprising sample RNA, adapter and handles, was performed in two separate hybridization steps, as described in Walbrun et al.(1). Biotin-functionalized handles were brought to reaction with the ssDNA adapter, while digoxigenin handles were directly attached to the sample RNA. Handle DNA was diluted to a final concentration of 0.071 µM and mixed with the molecule to anneal to in a 1:1 molar ratio in the hybridization buffer, then hybridized (for thermal protocols see **Tables S7** and **S8**). In a second hybridization reaction, the products of the first step were joined and incubated over night at room temperature. Final products were quality controlled *via* agarose gel electrophoresis (1.5%, TAE buffer), and stored at -80°C. In the end, the complete dumbbell construct is formed when the hybridized construct is connected to Digoxigenin and Streptavidin functionalized micron-sized silica beads that are trapped in a focused laser beam (**Fig. 1B**).

#### Optical trap measurements and data analysis

##### *Measurement Preparation*

To avoid simultaneous binding of multiple molecules to the same bead pair, assembled constructs were diluted to around 0.1 nM and incubated at room temperature for 10 min with micron-sized streptavidin-coated (SV) beads (Bangs Laboratories, Inc.) in HS buffer (20 mM $MgCl_2$, 300 mM KCl, 50 mM HEPES). After incubation, this solution was diluted into 300 µl of HS buffer. In similar manner, anti-digoxigenin-coated (AD) beads were diluted to identical concentration in HS buffer with the same scavenger system. In parallel, 500 µl of each running buffer for the three flow channels were prepared by addition of the scavenger system (final concentrations: 26 U/ml glucose oxidase (SIGMA-ALDRICH), 17 000 U/ml catalase (SERVA), and 0.65 % glucose (SIGMA-ALDRICH)) to the corresponding measurement buffer. A commercial microfluidic chip (C-Trap® Optical Tweezers - Fluorescence & Label-free Microscopy (LUMICKS)) was used that contains two bead channels and additionally allows maintenance of three different phases during measurement, separated by laminar flow (**Fig. S5**). For measurements with protein, one HS and one LS flow channel was usually run for initial quality control of the tether, while coreROQ or extROQ was added to LS buffer for the third channel in varying final concentrations (indicated correspondingly in the results). All five mixtures were injected into the syringe pump of the C-Trap® Optical Tweezers – Fluorescence & Label-free Microscopy. Laminar flow was obtained and diffusion between channels prevented by applying a pressure of ~ 0.35 bar to the syringes, resulting in a velocity of ~ 20 μm/s in the flow channels. In each of the two bead channels respectively, an SV bead, pre-incubated with the assembled sample construct, and an AD bead was captured into one of the trapping lasers. Alternatively, the two bead types could be co-injected into a single channel, in which case one of them can be identified by a fluorescent label (note that **Fig. S5** only displays this simplified setup). Experiments were started by establishing a construct tether between two beads in the HS channel and verifying correct single tether formation in a few stretch-relax cycles showing the fingerprint trace of the sample molecule. Then, the molecule was kept in the channel or moved to different channels according to the specific interest of the experiment. All measurements were performed using a trap stiffness between 0.25 pN/nm and 0.40 pN/nm, and at temperatures of ~ 25°C. Data was downsampled from the initial sampling rate of 78.125 kHz by a factor of 3 before analysis.

#### ***Constant velocity traces***

For the force (F) vs. extension (e) curves in **Fig. 1C, 3A, S1A, S1B and S2** we used the so-called constant velocity mode where the distance between the traps was first increased with a constant velocity until the molecule was fully unfolded and then decreased to allow the hairpin to refold. We used a velocity of 100 nm/s for **Fig. 1C, 3A, S1A, S2** and 200 nm/s for **Fig. S1B**. The traces provided the fingerprint of the corresponding molecules and allowed classification of intermediate states during the unfolding/refolding process.

Modeling polymer elasticity in constant velocity cycles:

States in the force vs. extension traces, where the RNA was folded, were fit with the extensible worm-like-chain (eWLC) model(2)

$$F_{eWLC}(e) = \frac{k_B T}{p_{dsDNA}}\left[\frac{1}{4}\left(1 - \frac{e}{L_{dsDNA,linker}} + \frac{F}{K}\right)^{-2} - \frac{1}{4} + \frac{e}{L_{dsDNA,linker}} - \frac{F}{K}\right]$$

with $k_B T$ the thermal energy, $p_{dsDNA}$ the dsDNA linker persistence length, $L_{dsDNA,linker}$ the dsDNA linker contour length and $K$ the elastic stretch modulus. When parts of the RNA were unfolded, the elastic behavior can be modeled by the eWLC in series with a standard WLC(3)

$$F_{WLC}(e) = \frac{k_B T}{p_{ssRNA}}\left[\frac{1}{4}\left(1 - \frac{e}{L_{ssRNA}}\right)^{-2} - \frac{1}{4} + \frac{e}{L_{ssRNA}}\right]$$

where $p_{ssRNA}$ is the persistence length of the unfolded ssRNA and $L_{ssRNA}$ the contour length of the unfolded ssRNA. For the fits with unfolded ssRNA ($F_{eWLC}$ in series with $F_{WLC}$) we used a fixed $L_{dsDNA,linker}$ , $p_{dsDNA}$ and $K$ from the previously fitted folded state of the same constant velocity trace. Also, the $p_{ssRNA}$ was fixed to 0.9 nm in the fitting process. Based on the increase in unfolded contour length from one to another state obtained in the WLC fits, the number of opened base-pairs could be calculated.

#### ***Passive-mode traces***

To get a more precise assignment of intermediate states, their lifetimes and to determine a detailed folding/refolding pathway, we performed so-called passive-mode experiments, where the distance between the lasers was held constant. This allowed observation of fluctuations of the RNA between its several intermediate states close to equilibrium in real time (see **Fig. 2B,**

**4F, S1C, S6, S9, S10A and S10C**). In the corresponding force vs. time traces states are more folded towards higher forces and more unfolded at lower forces as the folding process shortens the length of the construct, and hence, the beads are pulled out of the trap, increasing the load. Assignment of each data point to a certain state was done via Hidden-Markov-Modeling (HMM)(4). The assigned time-trajectory was then used to calculate the transition rates (see **Fig. S4A, S10E and S10F**)(5) and derive the folding network (see **Fig. 2D and S4B**).

The state-assigned traces were then used to extract free energies at zero load using a model previously introduced for protein folding under force that takes also the energetic contributions from the dumbbell assay into account(6). This model was adapted for RNA(1) and can be summarized as follows: the hybridized dumbbell construct bead-$DNA_{linker}$-RNA-adapter-$DNA_{linker}$-bead is first simplified to a bead-$DNA_{linker}$-RNA system with a combined contour length of both DNA-linkers (including the hybrid DNA adapter RNA overhang part), an effective trap stiffness $k_{eff}^{-1} = k_1^{-1} + k_2^{-1}$ and an effective bead deflection $x_{eff} = |x_1| + |x_2|$. In case of passive mode experiments, where the distance is held constant and not the force, each transition is accompanied by a change in extension of the DNA linkers and a change in the bead deflection. Therefore, the energetic contributions of these changes have to be considered in the zero force free energy calculation. The total free energy $G_i(F_i)$ stored in the system at force $F_i$ can be described by the individual contributions of the beads $G_i^{beads}(F_i)$, the stretched dsDNA linkers $G_i^{dsDNA,linker}(F_i)$, the unfolded ssRNA $G_i^{ssRNA}(F_i)$ and the folded RNA $G_{0,i}^{RNA}(F_i)$ in state *i*

$$G^{beads}(F) = \frac{1}{2} k_{eff}^{-1} F^2$$

$$G^{dsDNA,linker}(F) = \int_0^{e_{eWLC}(F)} F_{eWLC}(e')de'$$

$$G^{ssRNA}(F) = \int_0^{e_{WLC}(F)} F_{WLC}(e')de'.$$

The parameters of the eWLC and WLC were obtained from previous fits to the force-extension traces explained in the previous section. By using the Boltzmann equation, the free energy difference between two states *i,j* at forces $F_i$ and $F_j$ can be calcultated as

$$\Delta G_{ij}(F_i, F_j) = G_j(F_j) - G_i(F_i) = -k_B T \cdot ln\left(\frac{P_j(F_j)}{P_i(F_i)}\right).$$

Combining this with the previously defined individual contributions, the free energy difference between two RNA intermediate states at zero force is then

$$\Delta G_{0,ij} = -k_B T \cdot ln\left(\frac{P_j(F_j)}{P_i(F_i)}\right) - \Delta G^{beads}(F_i, F_j) - \Delta G^{dsDNA,linker}(F_i, F_j) - \Delta G^{ssRNA}(F_i, F_j).$$

For the probabilities $P_i(F_i)$ and $P_j(F_j)$, we used the HMM assigned state population probabilities.

To extrapolate transition rates to zero load (**Fig. S10 E and F**), we repeated the passive mode experiments at different distances (**Fig. S9**) and therefore obtained transition rates $k_{ij}(F_i)$ at different forces $F_i$. To model the force dependence of the transiton rates, we use the contour length increase during an unfolding/refolding transition as a reaction coordinate that is well defined as RNA unfolding/refolding is characterized by step by step zipping/unzipping of the base pairs. Note that a simple Bell model that assumes a force independent distance to the transition state is not applicable here. The effective length to reach the transition state depends on the extension of the unfolded ssRNA which is highly force dependent and can be described by a WLC. Also, the contributions of the beads and the dsDNA-linkers have to be considered. Here, we use a model first introduced for unfolding/refolding of globular proteins(7) and coiled coils(6) and later adapted for RNA(1). To summarize: We define the free energy barrier $\Delta G_{iT}^{\#}$ from the initial state *i* to the transition state *T* that takes all the energy changes of the dsDNA linkers, springs and unfolded ssRNA into account

$$\Delta G_{iT}^{\#}(F_i, F_T) = \Delta G_{iT}^{beads}(F_i, F_T) + \Delta G_{iT}^{dsDNA,linkers}(F_i, F_T) + \Delta G_{iT}^{ssDNA/ssRNA}(F_i, F_T)$$

with $F_i$ the force that acts on the molecule in state *i* and $F_T$ the force that acts on the molecule at the transition state. Again, the WLC fitting parameters from the force-extension traces are used in the calculation of the free energies of the dsDNA linkers and the unfolded ssRNA. The force dependent transition rates can then be calculated as

$$k_{ij}(F) = k_{0,ij}\exp\left(-\Delta G_{iT}^{\#}(F_i = F, F_T)/k_B T\right)$$

with $k_{0,ij}$ the folding rate constant at zero load. The only free fitting parameters are then $k_{0,i}$ (y intercept in **Fig. S10 E and F**) and the contour length difference $\Delta L_{iT}^{\#}$ from state *i* to the transition state *T* that defines the slope of the fits at infinite force.

#### *Off-rates and on-rates of Roquin-binding*

To obtain the off-rates of coreROQ or extROQ binding to the isolated ADE element, we used the HMM assigned passive mode traces. As shown schematically in **Fig. S7**, the ROQ bound states are characterized by an increased lifetime of I3. Since the lifetime of the folded not bound and folded bound state differs over two orders of magnitude, the population of Roquin-bound events is easily separable. The off-rate $k_{off}$ is then calculated as the inverse of the average lifetime $\langle\tau_{bound}\rangle$ of the Roquin bound states:

$$k_{off} = \frac{1}{\langle\tau_{bound}\rangle}$$

The off-rates show a force dependence that hints at an additional unbinding pathway, e.g. from the folded bound state to the unfolded bound state to the unfolded unbound state. The off-rates can then be explained by a force dependent and a force independent rate contribution

$$k_{off}(F) = k_{off,0,0} + k_{off,0,1} \cdot exp(b \cdot F)$$

with $k_{off,0,0}$, the force-independent contribution to the off-rate, and parameters $k_{off,0,1}$ and $b$ defining how strong force couples to the off-rate. By fitting the off-rates shown in **Fig. 3B**, the off-rate at zero load was determined as $k_{off,0} = k_{off,0,0} + k_{off,0,1}$. The values are summarized in **Table S3**.

The on-rate $k_{on}$ is calculated via the inverse of the average lifetime of the Roquin unbound states and divided by the Roquin concentration:

$$k_{on} = \frac{1}{\langle\tau_{unbound}\rangle \cdot [R]}$$

Since the on-rate of Roquin to the unfolded compared to the not-unfolded state differs, $k_{on}$ decreases with increasing force (see gray data points in **Fig. S7C**), where the molecule is more often in the unfolded state. To calculate the on-rate to the not-unfolded state, meaning every state folded to at least I3, we have to adapt the on-rate to

$$k_{on,\overline{U}} = \frac{1}{\langle\tau_{unbound}\rangle \cdot \frac{P_{\overline{U},unbound}}{P_{unbound}} \cdot [R]}$$

with $P_{\overline{U},unbound}$ the probability to be in the not-unfolded and unbound state and $P_{unbound}$ the probability to be in the unbound state. The on-rate $k_{on,\overline{U}}$ no longer shows a significant force dependence (**Fig. S7C**) and the on-rate at zero load $k_{on,\overline{U},0}$ can simply be calculated as the average of $k_{on,\overline{U}}$. The $K_D$ at zero load was then determined as

$$K_{D,0} = \frac{k_{off,0}}{k_{on,\overline{U},0}}.$$

### SUPPLEMENTARY TABLES

The standard error of the mean (s.e.m.) is indicated in brackets behind a mean value, if not stated otherwise. Its value is calculated using a two-sided student-t distribution with a confidence interval of 68.2 %.

**Table S1.** Cumulated gains in unfolded contour length by unfolding/refolding transitions of *Ox40* 3'UTR, derived from WLC fits to stretch-relax cycles of FL (N = 5) and its sub-constructs CDE (N = 11), ADE (N = 24) and ADE-CDE (N = 9). The expected contour length is calculated using a conversion rate of 0.6 nm/nt(1). As the transitions downstream of the Terminus region occur simultaneously and overlap in fl, they are subsumed under the "Total" transition up to the unfolded state.

| Construct | Transition | Expected Contour Length [nm] | Measured Contour Length [nm] |
|---|---|---|---|
| CDE | $F_{CDE} \leftrightarrow U_{CDE}$ | 7.8 | 7.78 (0.23) |
| ADE | F ↔ I1 | 6.6 | 7.33 (0.15) |
| | I1 ↔ U | 28.2 | 28.19 (0.19) |
| ADE-CDE | F ↔ I1 | 6.6 | 7.17 (0.33) |
| | $F_{CDE} \leftrightarrow U_{CDE}$ | 14.4 | 14.58 (0.36) |
| | I1 ↔ U | 36.0 | 35.85 (0.49) |
| FL | $\text{Terminus}_{\text{default/alt}}$ | 34.8/29.4 | 31.03 (0.59) |
| | Total | 82.8 | 82.31 (1.08) |

**Table S2.** Relative free energies of states making up the ADE unfolding/refolding landscape, extracted from passive mode experiments in HS buffer via population probabilities. Values indicate the difference in free energy between two folds, calculated for zero-force. Data from three different force biases was used for U-I3 and I3-I2, while four different forces could be taken into account for I2-I1, I1-M and M-F. Note that the value for U-I3 has a higher uncertainty due to the unfolded state's undermost position on the force axis, leading to error-prone assignment of data points at low force. In total, the obtained free energies amount to an energy of -25.0 kcal $mol^{-1}$ for natively folded ADE.

| Interval | U-I3 | I3-I2 | I2-I1 | I1-M | M-F |
|---|---|---|---|---|---|
| $\Delta G_0 [\text{kcal mol}^{-1}]$ | -7.99 (2.16) | -5.83 (0.27) | -7.31 (0.43) | -1.75 (0.15) | -2.16 (0.27) |

**Table S3.** On- and off-rates of coreROQ and extROQ to ADE. The on- and off-rates from Fig. 3B, C were fit to a model described in the SI. We iteratively adapted the parameters $k_{off,0,1}$ and $b$ that define how strong force couples to the off-rate to obtain maximal overlap between data and model and then only used $k_{off,0,0}$, the force-independent contribution to the off-rate, as free fitting parameter. The value in the brackets for $k_{on,\overline{U},0}$ represents the s.e.m. and for $k_{off,0,0}$ the 1σ standard deviation of the fitting parameter. ADE + coreROQ (N = 27) and ADE + extROQ (N = 23).

| System | $k_{on,\overline{U},0}$ [1/(s µM)] | $k_{off,0,0}$ [1/s] | $k_{off,0,1}$[1/(1000 s)] | *b* [1/pN] |
|---|---|---|---|---|
| ADE + coreROQ | 241 (22) | 1.15 (0.09) | 0.9 | 0.6 |
| ADE + extROQ | 292 (28) | 0.30 (0.03) | 0.1 | 0.6 |

**Table S4.** Folding and unfolding transition rates from ADE I1 to I3. The transition rates from **Fig. S10 E and F** were fit to a model described in the SI “Passive mode traces”. The contour length differences $\Delta L^{\#}_{I1,T}$ from state I1 to the transition state, $\Delta L^{\#}_{T,I3}$ from the transition state to state I3 and the zero-force unfolding rate $k_{0,I1,I3}$ and refolding rate $k_{0,I3,I1}$ were gained from the fit for the three systems ADE, ADE with coreROQ bound and ADE with extROQ bound. The value in the brackets represent the 1σ standard deviation of the fitting parameter. ADE (N = 15), ADE + coreROQ (N = 11) and ADE + extROQ (N = 9).

| System | $\Delta L^{\#}_{I1,T}$ [nm] | $\Delta L^{\#}_{T,I3}$ [nm] | $k_{0,I1,I3}$[1/s] | $k_{0,I3,I1}$ [$10^5$/s] |
|---|---|---|---|---|
| ADE | 5.5 (5.2) | 7.6 (2.6) | 0.08 (0.31) | 2.9 (5.0) |
| ADE + coreROQ | 5.4 (3.1) | 7.5 (3.6) | 0.10 (0.25) | 1.53 (4.2) |
| ADE + extROQ | 4.5 (1.2) | 8.4 (1.4) | 0.32 (0.27) | 0.87 (0.88) |

**Table S5.** Sequences for optical tweezer experiments. The primers Fw_primer_dig, Fw_primer_bio and Bw_primer were used in the PCR reaction for the dsDNA linker preparation. The [DIG] and [BIO] insertion shows where the biotin and dig-oxigenin modifications are placed. The X in the Bw_primer stands for the abasic site. Lambda-phage DNA was used as a template, and the forward and backward primers were designed to amplify the region containing base pairs 10557-11100.

| Name | Sequence (5’-3’) |
|---|---|
| Fw_primer_dig | [DIG]CA GAG CGC CA[DIGdT] TG[DIGdT] GAA GGC G |
| Fw_primer_bio | [BIO]CA GAG CGC CA[BIOdT] TG[BIOdT] GAA GGC G |
| Bw_primer | CCT CCC GCA CCA CTC CAA CTA CGT CAG CCC TGC CXG CGG GCT TCC TGA AAA CTG A |
| Adapter | GGC AGG GCT GAC GTA GTT GGA GTG GTG CGG GAG GCA CCT GAA TTC TGC CCT CAC T |
| RNA 5’-overhang | GGC AGG GCU GAC GUA GUU GGA GUG GUG CGG GAG G |
| RNA 3’-overhang | AGU GAG GGC AGA AUU CAG GUG |
| CDE | 5’-overhang-GGC UUU CCU AUG UAU GCU AUG CAU ACU AC-3‘-overhang |
| ADE | 5’-overhang-GCC UGC CAG UAC CCU CCA CAC CGU UCU AGG UGC UGG GCU GGC UCU GGG C-3‘-overhang |
| ADE-CDE | 5’-overhang-GCC UGC CAG UAC CCU CCA CAC CGU UCU AGG UGC UGG GCU GGC UCU GGG CUU UCC UAU GUA UGC UAU GCA UAC UAC-3‘-overhang |
| fl | 5’-overhang-GCA UUA CUA CAG GAG UGG AUU UUA UGG GGC ACG GAC AAC CCA UAU CCU GAU GCC UGC CAG UAC CCU CCA CAC CGU UCU AGG UGC UGG GCU GGC UCU GGG CUU UCC UAU GUA UGC UAU GCA UAC UAC CUG CCU GGU GGU GC-3‘-overhang |

**Table S6.** Thermal conditions of the polymerase chain reaction (PCR).

| Time | Temperature |
|---|---|
| 5 min | 94°C |
| 15 s<br>15 s<br>40 s | 94°C<br>63°C 45 x<br>68°C |
| 5 min | 68°C |
| stored at 4°C | |

**Table S7.** Thermal conditions applied during hybridization of biotin handles and adapter.

| Time | Temperature |
|---|---|
| 10 min | 75°C |
| 10 min | 65°C |
| 10 min | 55°C |
| 10 min | 45°C |
| 10 min | 40°C |
| 10 min | 35°C |
| 10 min | 30°C |
| 10 min | 25°C |
| 1 min | 20°C |
| 1 min | 15°C |
| Stored @ 4°C | |

**Table S8.** Thermal conditions applied during hybridization of digoxigenin handles and sample RNA.

| Time | Temperature |
|---|---|
| 10 min | 80°C |
| 40 min | 62°C |
| 40 min | 52°C |
| 1 min | 45°C |
| 1 min | 40°C |
| 1 min | 35°C |
| 1 min | 30°C |
| 1 min | 25°C |
| 1 min | 20°C |
| 1 min | 15°C |
| Stored @ 4°C | |

### SUPPLMENTARY FIGURES

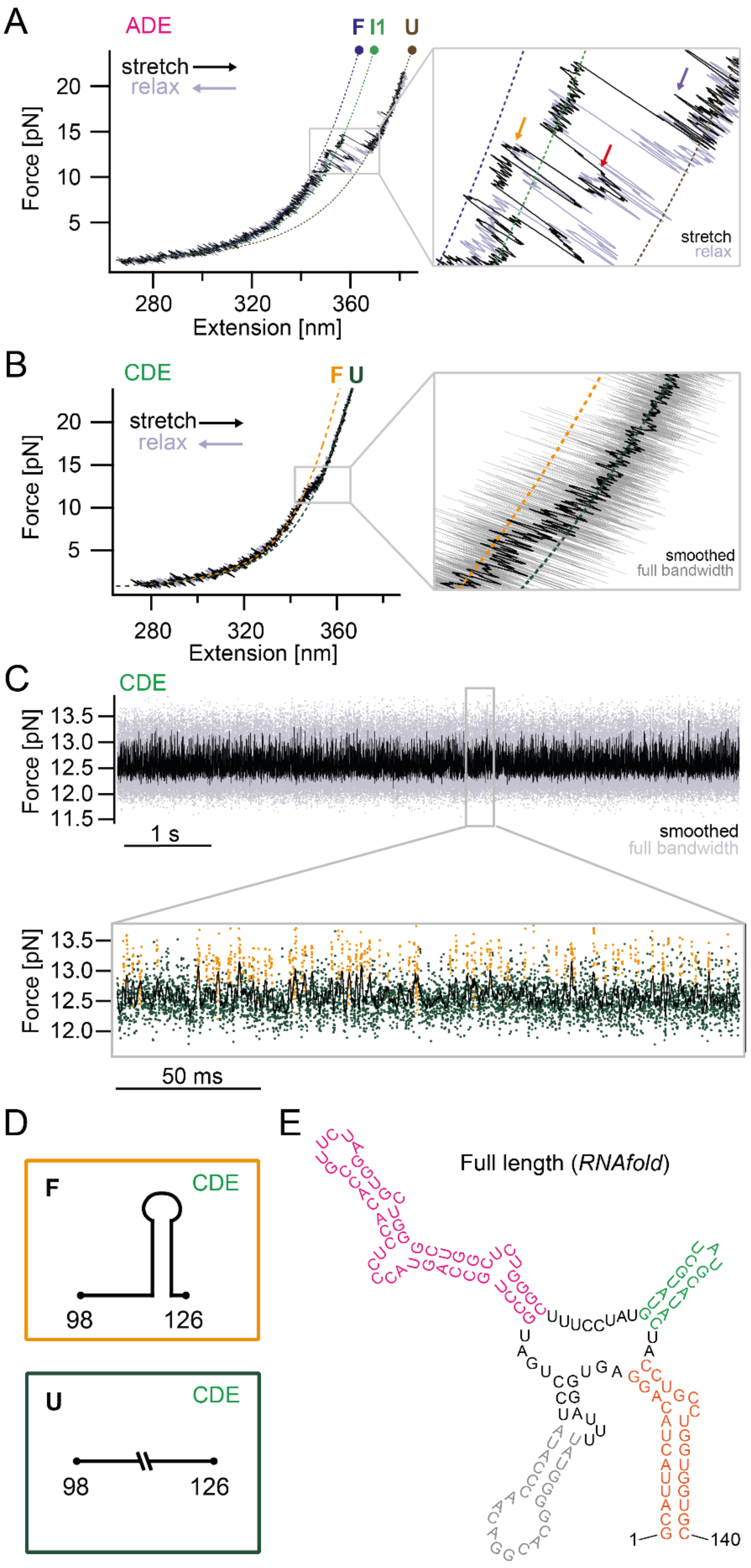


**Figure S1.** In-depth characterization of the *Ox40* ADE and CDE. A) Individual force-extension curve of ADE, featuring multiple intermediates. WLC fits were applied to the three states of this pathway F (folded, dark blue), I1 (intermediate 1, green), and U (unfolded, brown). A zoom into the trace magnifies the transition towards the unfolded state and exposes several short-lived states (arrows in orange, red, purple). B) Stretch-relax cycle of the CDE element, pulled at 200 nm/s in high salt (HS) buffer. The CDE folds as one well-defined stem-loop and exhibits a single, but highly dynamic transition from the folded (F, orange) to the unfolded (U, green) state. The zoom shows smoothed (black) and full-bandwidth (gray) data of the stretching curve, demonstrating how the fast transition leads to a hump-like shape in the smoothed curve. C) Passive mode experiment of CDE in HS buffer. The upper trace shows multiple seconds of the CDE flipping between its folded and unfolded states. A zoom into a ≈200 ms excerpt (lower) discloses individual folding and unfolding events. D) Schematic depictions of the folded (F) and unfolded (U) state of CDE. E) Secondary structure prediction of full-length *Ox40* 3'UTR showing the additional base pairing adjacently to the Bulge using *Vienna RNAfold(8)*.

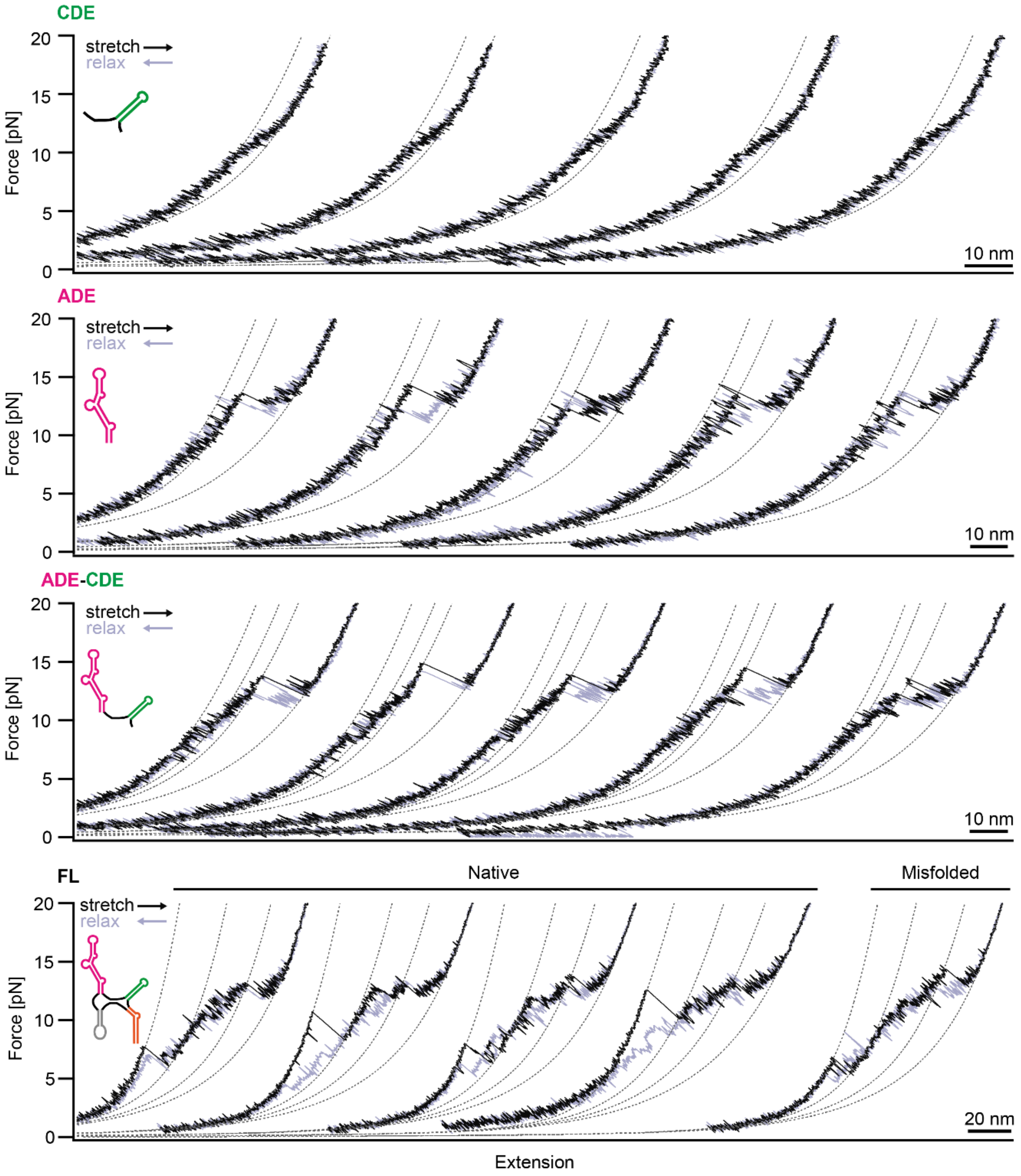


**Figure S2.** Representative force-extension curves of examined RNA constructs in HS buffer. WLC fits were applied as described in the main text (dashed lines). All traces are recorded at a pulling velocity of 100 nm/s. The CDE shows reproducible unfolding/refolding transitions close to equilibrium. The two main transitions of the ADE construct exhibit slower kinetics compared to the CDE and the traces show a certain hysteresis. Note that for ADE-CDE, the single unfolding transition of CDE overlaps with and thereby distorts the F-I1 transition of the ADE. The full-length construct comprising the entire *Ox40* 3'UTR except its unstructured 3' tail frequently appears to engage in non-native refolding, resulting in deviating unfolding traces. One of these traces, which can be discerned by an insufficient total unfolded contour length, is shown on the right, while the first four traces represent natively unfolding fl construct. Comparison of the WLC fits indicates that an incorrectly folded Terminus, resulting in a lower distance between the 1$^{st}$ and 2$^{nd}$ fit, mainly accounts for incorrect folding. This reflects in a lower average unfolded contour length value of 24.5 ± 0.3 nm for misfolded Terminus transitions, deviating from native folding transitions (31.0 ± 0.6 nm) by 6.5 nm, while the total unfolded contour length of 77.3 ± 0.5 nm deviates from correct total unfolding (82.3 ± 1.1 nm) by 5.0 nm.

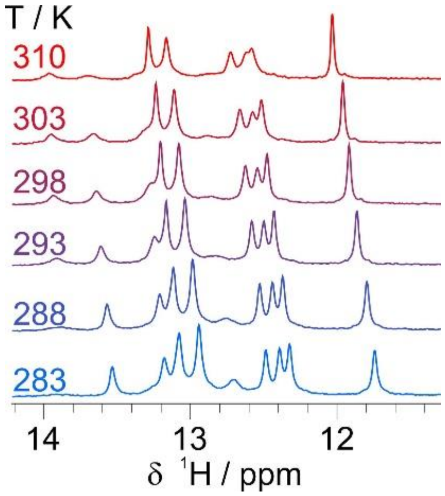


**Figure S3.** $^{1}H$ imino spectra of apo ADE at varying temperatures.

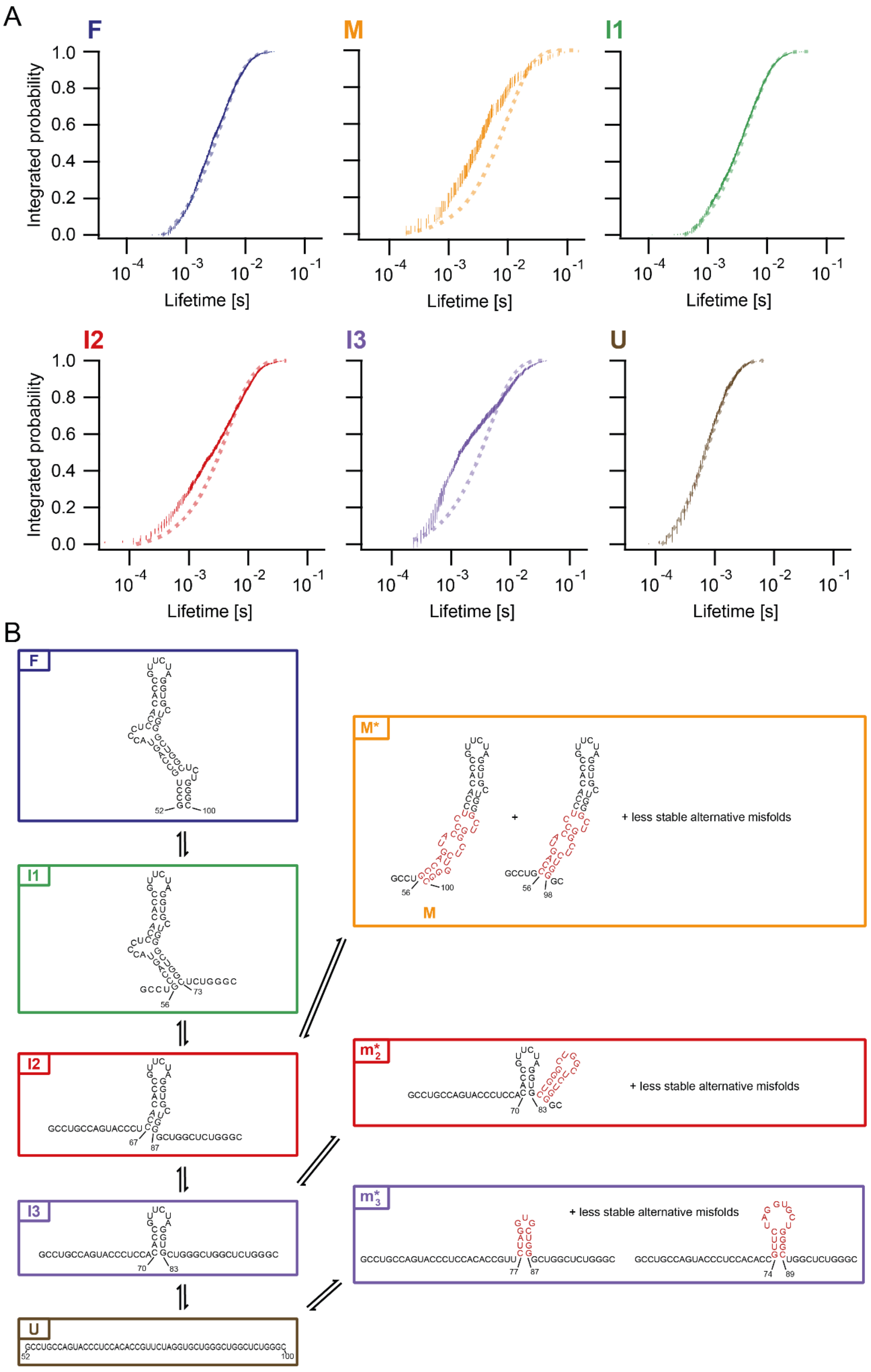


**Figure S4.** Folding network of the ADE. A) Lifetime distributions of states constituting the ADE folding/unfolding network, extracted from passive mode experiments. Cumulative relative frequencies of lifetimes are plotted semi-exponentially against the lifetimes. While F, I1 and U display single-exponential distributions, lifetimes of I3, I2 and M deviate from a single-exponential fit and are thus distributed multi-exponentially. F recorded at 10.0 pN (N = 2414), M at 14.5 pN (N = 179), I1 at 9.5 pN (N = 2427), I2 at 13.3 pN (N = 1687), I3 at 12.8 pN (N = 911) and U at 12.2 pN (N = 697). Error bars are computed via Bootstrapping with a 68.2 % confidence interval. B) Structural model for the folding/unfolding network of ADE. States of the same color correspond to identical HMM states, i.e. states with similar or identical number of unfolded nucleotides. Ensembles of functionally redundant states on the same force level are highlighted with asterisks *. Non-natively folded nucleotides are indicated in red. The pathway of ADE features three on-pathway intermediates (I1, green; I2, red; I3, purple) between the folded (F, blue) and the unfolded (U, brown) state. Moreover, there are two off-pathway misfolded state ensembles ($m_{I2}$*, red; $m_{I3}$*, purple) on the levels I2 and I3 as well as an off-pathway misfolded state ensemble populating a unique force level between I1 and F (M*, orange). Representative structure candidates conceived with the help of the *RNAstructure fold*(9*)* server and *Nupack*(10) are indicated for $m_{I3}$* and $m_{I2}$*. To determine the likeliest candidate for the main misfold structure M, the relative position of this state on the force axis together with predictions from *RNAstructure fold*(9) were used. (full ADE sequence, max. energy difference = 50 %, max. number of structures = 25, window size = 0, maximum loop size = 10, temperature = 298 K).

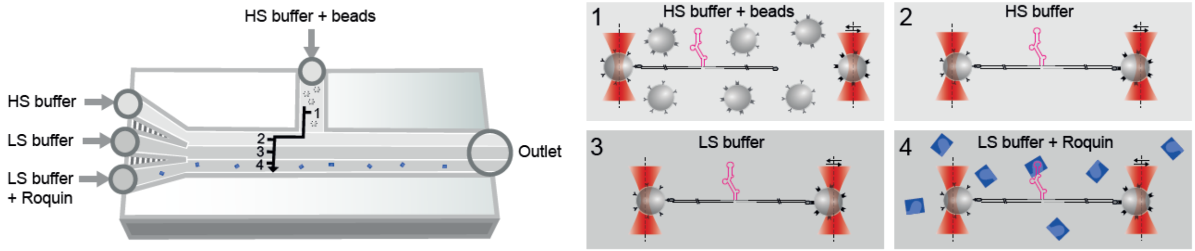


**Figure S5.** Spatial organization and measurement principle of the microfluidic chamber used for dual-beam optical tweezers experiments. Streptavidin-coated beads are pre-incubated with the assembled construct comprising sample RNA, ssDNA adapter and dsDNA handles. Together with anti-digoxigenin coated beads, they are flushed into the bead channel (1), where each of the bead types is trapped into one of the optical traps. The traps are then moved to the flow channel filled with stabilizing high salt (20 mM $MgCl_2$, 300 mM KCl, 50 mM HEPES) (HS) buffer (2), where they are closely approached to allow tether formation. Correct tether formation is usually verified by a few stretch-relax cycles showing the characteristic fingerprint trace of the molecule. From here, the same molecule can be subjected to further stretch-relax cycles or passive mode experiments in different conditions, such as in low salt (150 mM NaCl, 20 mM Tris) (LS) buffer (3) or with addition of core/extROQ (4), which require LS buffer for binding.

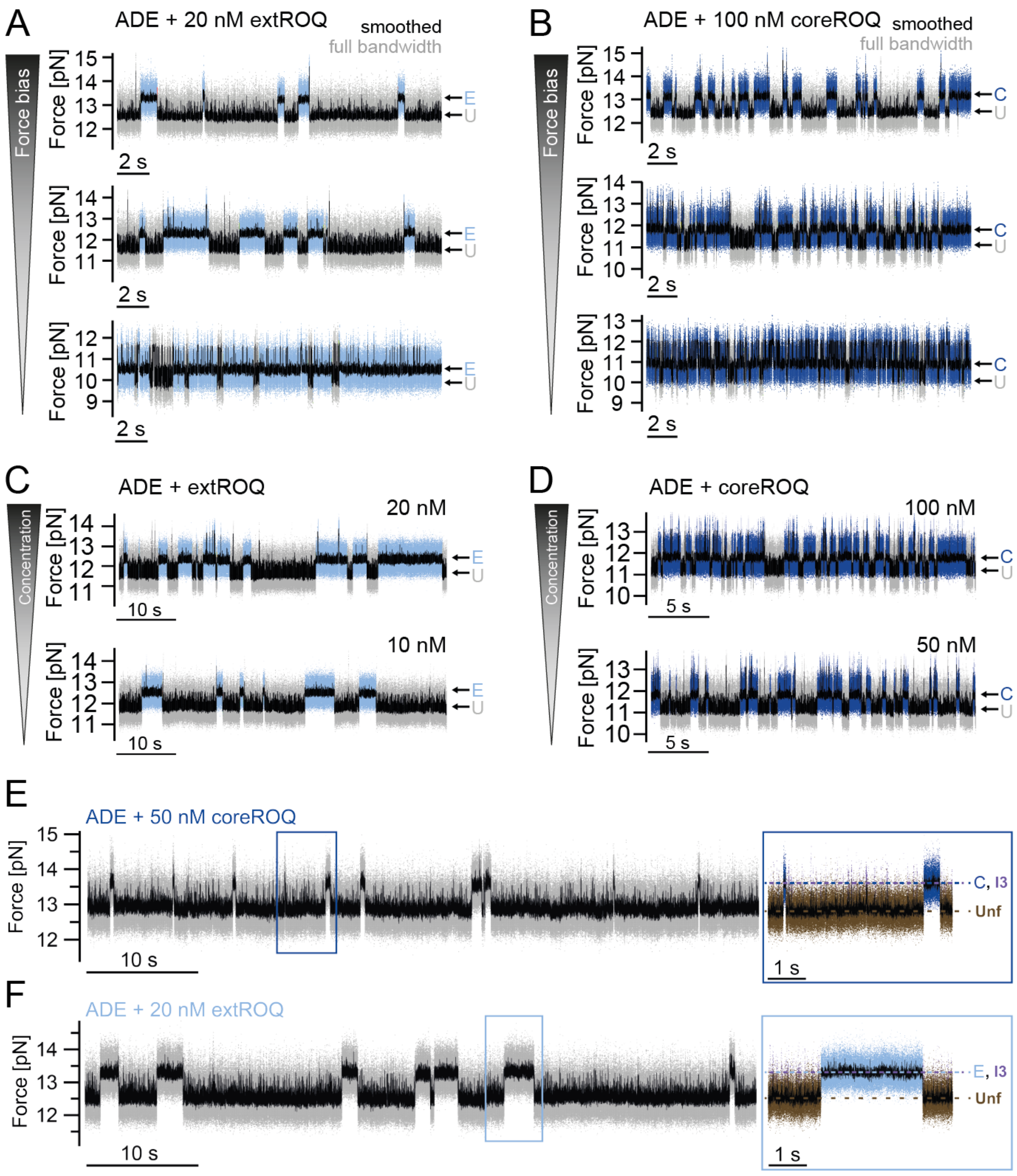


**Figure S6.** Passive mode traces of ADE with ROQ variants under differing conditions. Since coreROQ exhibits weaker binding than extROQ, experiments with coreROQ were generally performed at higher concentrations. A) 20 nM extROQ, descending force bias from upper to lower traces. B) 100 nM coreROQ, descending force bias from upper to lower traces. C) 20 and 10 nM extROQ at similar forces. D) 100 and 50 nM coreROQ at similar forces. E) 50 nM coreROQ. F) 20 nM extROQ. As the force decreases, the apical stem-loop is populated more frequently, offering more binding opportunities to extROQ (A) or coreROQ (B). Additionally, unbinding rates decrease for both proteins with descending force levels, which can be recognized by increased lifetimes of bound states. This can be explained by a force-dependent contribution to off-rates, i.e. the presence of a partially accessible force-dependent unbinding mechanism in addition to the force-independent unbinding followed by unfolding of the ADE. The zooms of E) and F) display representative parts of the traces, HMM-assigned to the states U (unfolded, brown), I3 (intermediate 3, purple) and C (coreROQ bound, dark blue) or E (extROQ bound, light blue). They demonstrate the bound states to be on the same force level as I3, providing visual evidence for the apical stem-loop as the core/extROQ-stabilized target structure of ADE.

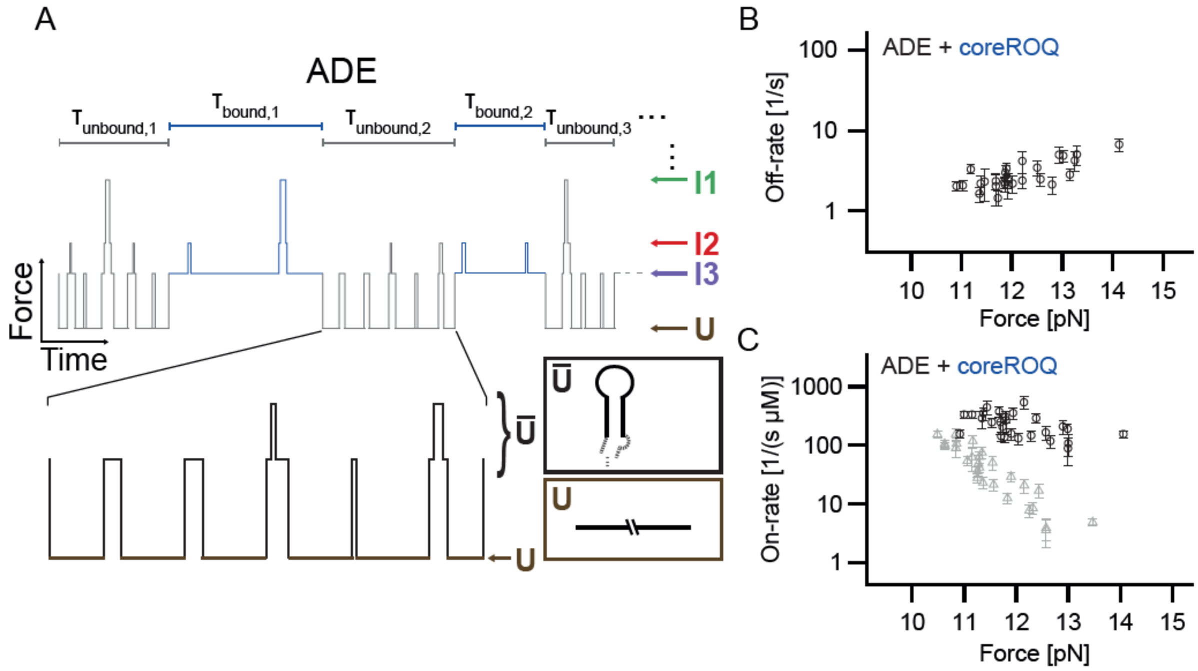


**Figure S7.** Calculation of on-/off-rates. A) Theoretical model for the extraction of on- and off-rates of ROQ proteins to ADE from passive mode traces. A schematic passive mode trace is shown with different force levels corresponding to different intermediates shown on the right. When ROQ proteins are bound to ADE, I3 is stabilized visible in prolonged lifetimes of I3 and lower unfolding rates from I3 to U. The bound phases are marked in blue, while the unbound phases are highlighted in gray. The off-rates can be calculated via the inverse of the average lifetimes of the bound states $\langle \tau_{bound} \rangle$. For the on-rates, one could take the inverse of the average lifetimes of the unbound states $\langle \tau_{unbound} \rangle$ and divide by the Roquin concentration. The unbound phases are a mixture of different states, however, where the ratio of different states highly depends on the applied force. If Roquin has a different binding kinetic e.g. to the unfolded state compared to I3, the observed on-rates will drastically change with force. Therefore, an effective on-rate is calculated considering the population probability ratio of unfolded (U) to all non-unfolded ($\overline{U}$) states (see SI "Off-rates and on-rates of Roquin binding" for more details). B) Off-rates of coreROQ from ADE. The force dependence reveals a force-dependent and force-independent contribution to the off-rate (see SI "Off-rates and on-rates of Roquin binding" for more details). The off-rates are the ones shown in Fig. 3B (N = 27). C) On-rates of coreROQ to ADE. The data points calculated with the non-effective method explained above are shown in gray (N = 27) and with the effective on-rate in black (N = 27). Using the non-effective method, the on-rate decreases significantly with force, which means that Roquin has a higher binding rate to folded compared to unfolded conformations. The effective on-rate is largely unaffected by force, which can be interpreted such that there is no significant difference in the binding kinetic of Roquin to the partially folded I3 compared to more folded structures.

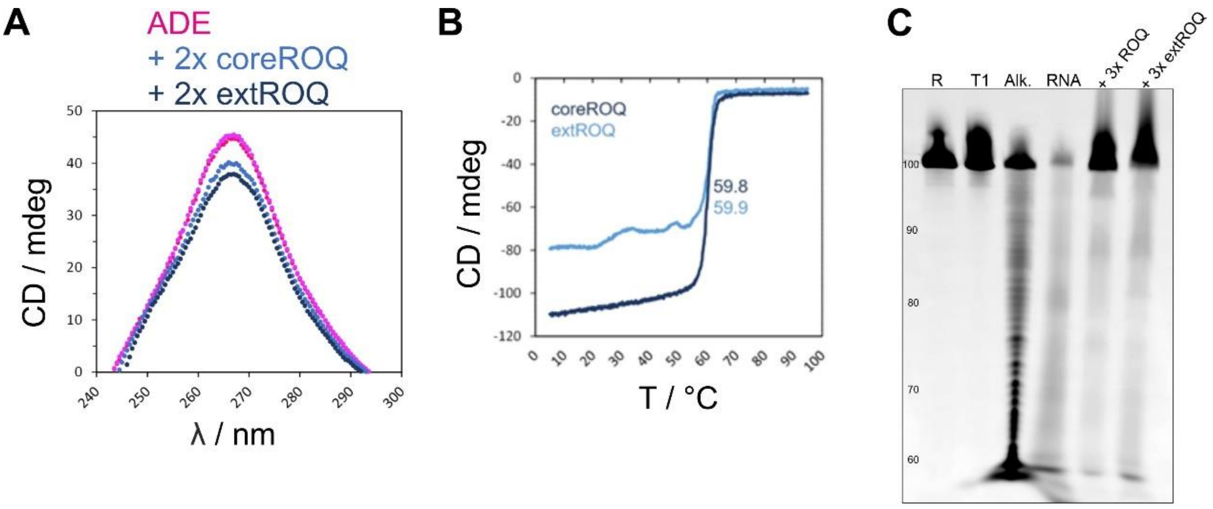


**Figure S8.** Roquin destabilizes ADE stem-loop structures. A) CD spectra of apo ADE (magenta) and in presence of core (dark blue) and extROQ (light blue). B) CD melting curves of core and extROQ. C) PAGE of OH• probing of apo ADE and in presence of coreROQ or extROQ.

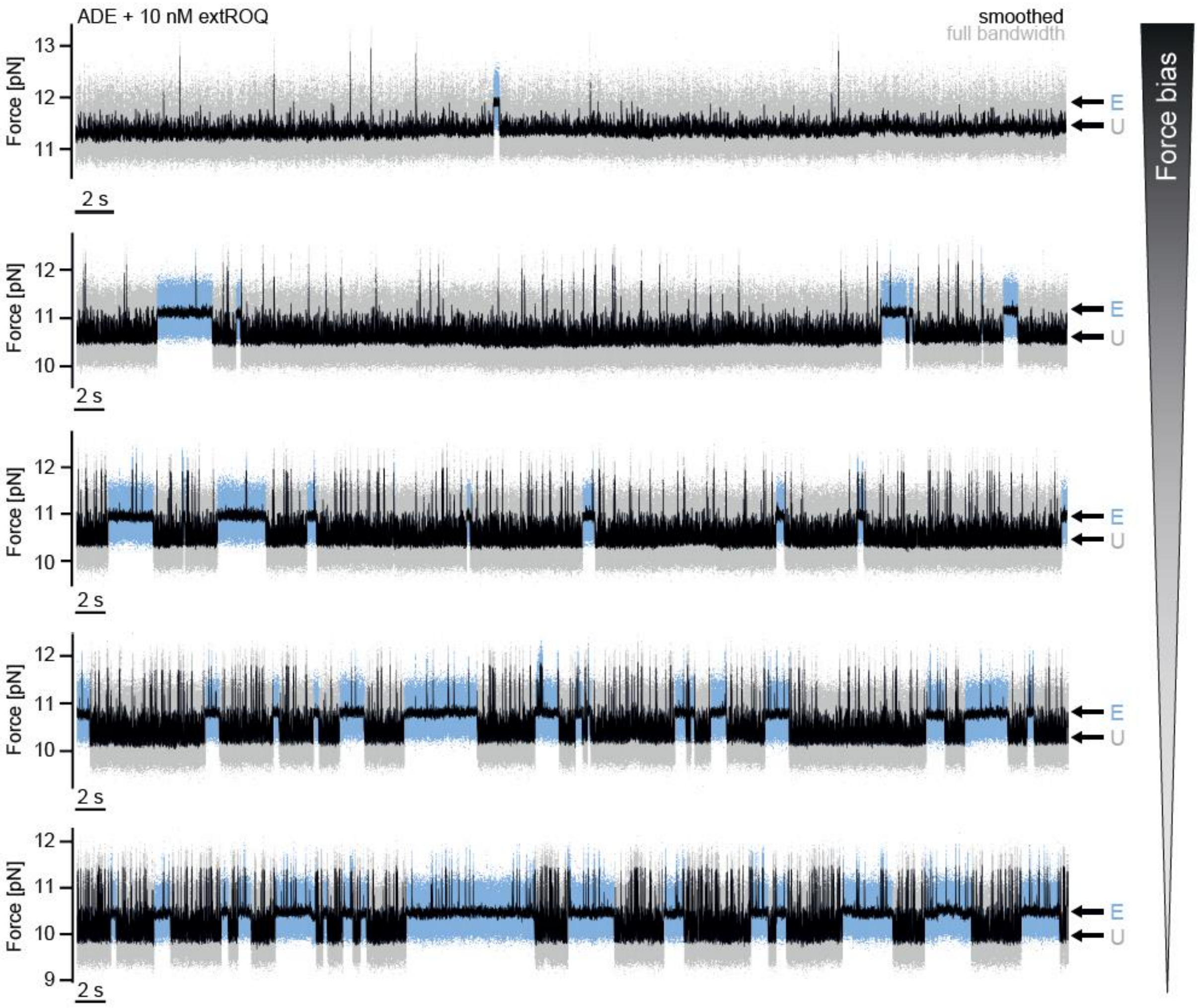


**Figure S9.** Passive mode experiments of ADE with 10 nM extROQ under decreasing force bias. At high force, the apo RNA transiently forms the apical I3 stem-loop, quickly unfolding again. Binding of extROQ stabilizes this hairpin structure (bound, E), increasing its lifetime. With decreasing force, the RNA can populate folded states more often, increasing the density of binding events by giving more binding opportunities to the protein. The lifetimes of bound states also increase, demonstrating the previously discussed force-dependence of the off-rate. Furthermore, especially at lower forces, the bound phases feature lower population of states folded further beyond I3 compared to the unbound phases. This points towards a destabilization of higher folds by extROQ, while it stabilizes the specific apical stem-loop structure formed by I3.

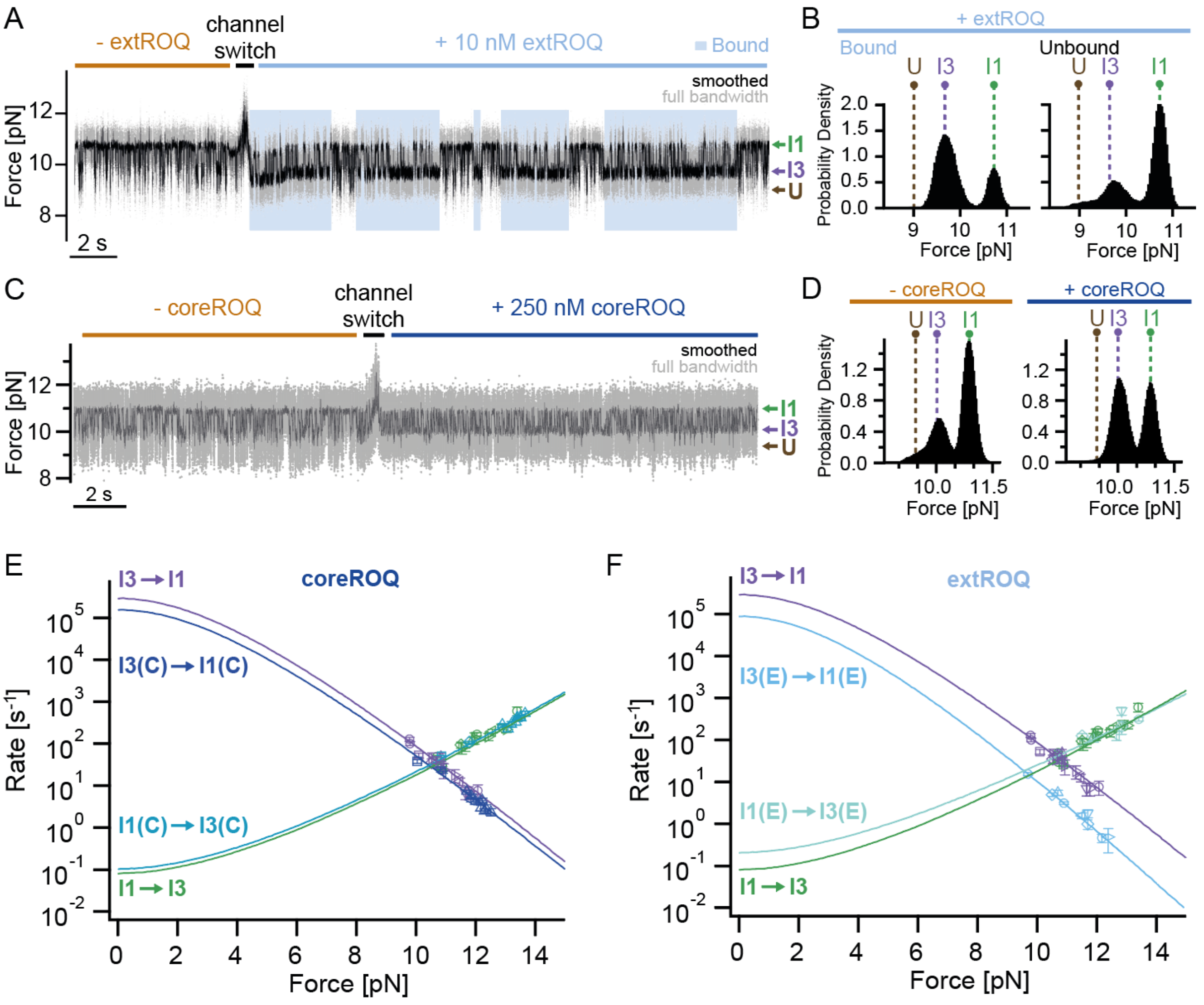


**Figure S10.** Destabilization of ADE secondary structure by coreROQ and extROQ. A) Passive mode trace of ADE in apo form (without extROQ, left) and in presence of 10 nM extROQ (right, bound phases are highlighted in light blue). B) Population distributions of ADE folds within the bound (left) and unbound (right) states of ADE during measurement with 10 nM extROQ. The histograms display the distribution of data points at different forces corresponding to the different states U, I3 and I1 (indicated in dashed lines). Stabilization of I3 by extROQ can be seen to come along with destabilization of I1. Note that data points of the short-lived intermediate I2 overlap with its neighboring I3 state, causing the latter to have an asymmetric skew towards higher forces in the unbound state. As the extROQ bound I3 state displays a much more symmetric, though still slightly right-bended distribution, the protein can be suspected to already destabilize more folded secondary structure at I2 level. C) Analogous passive mode trace with 250 nM coreROQ. D) Analogous histogram with coreROQ. Instead of the protein bound phase, the whole trace in the coreROQ channel is used for the histogram as ADE is almost exclusively bound by coreROQ at this concentration. E) Unfolding/refolding rates of the I1/I3 transition of coreROQ bound and unbound ADE. Rates were fitted using a rate model explained in SI "Passive mode traces" (see **Table S4** for fitting parameters). The unfolding rate from I1 to I3 is largely indifferent to the presence (light blue, ascending curve, N = 11) or absence (green, ascending curve, N = 15) of coreROQ, while the refolding rate from I3 to I1 is slightly slowed down by coreROQ (dark blue, descending curve, N = 11) in comparison to apo ADE (purple, descending curve, N = 15). F) Analogous rate plot with extROQ. Note that the refolding rate (light blue) is significantly reduced in the presence of extROQ. Refolding (extROQ, light blue, N = 9), refolding (apo ADE, purple, N = 15), unfolding (extROQ, turquoise, N = 9), unfolding (apo ADE, green, N = 15). Error bars represent the s.e.m..

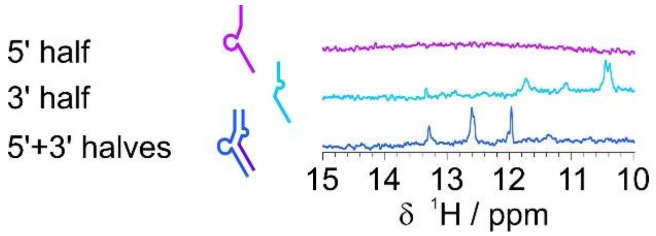


**Figure S11.** Structural integrity of ADE RNA fragments. $^1$H imino spectra of 5’ and 3’ ADE halves and the annealed duplex confirm ss and ds structures.

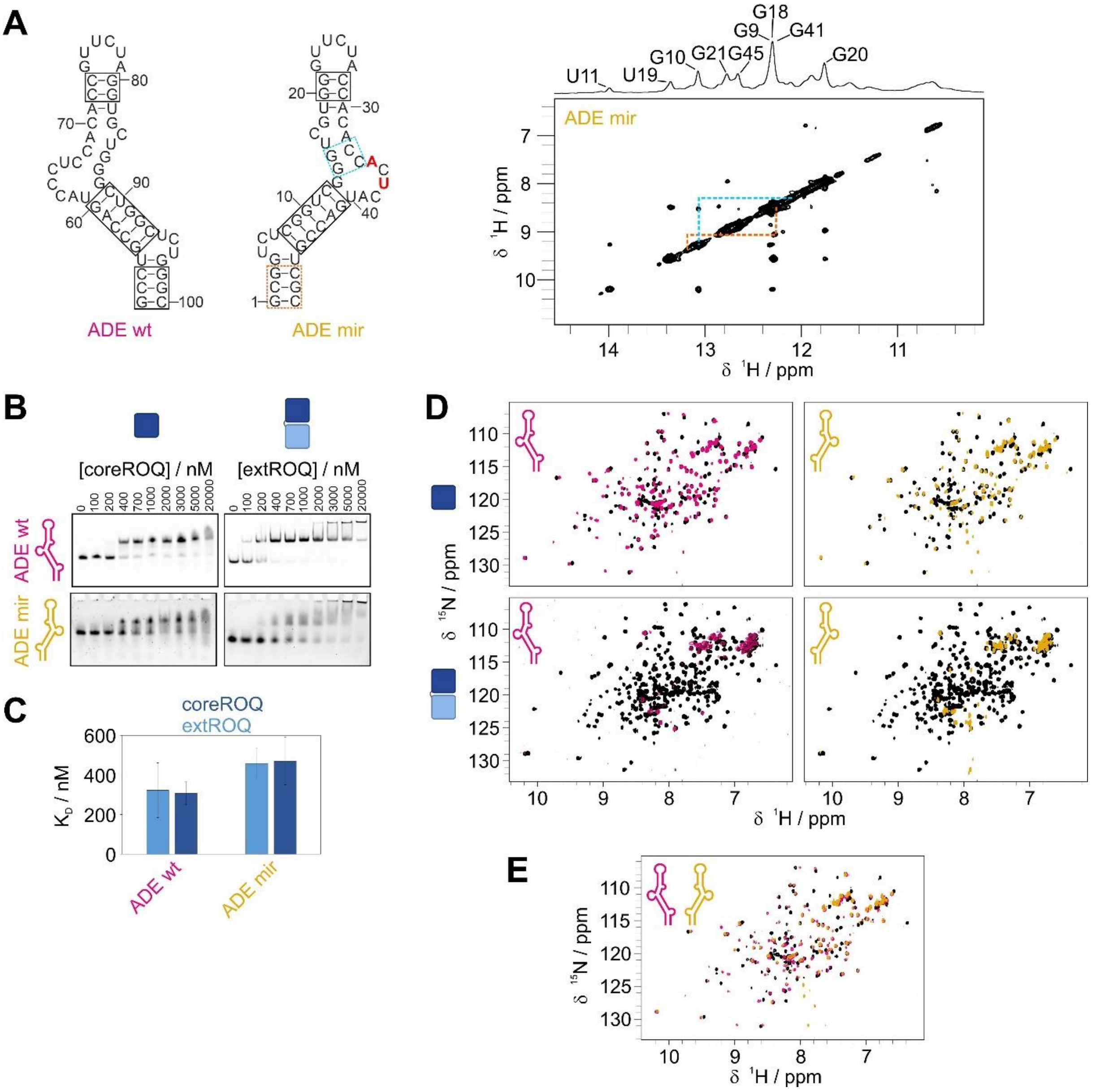


**Figure S12.** Roquin binding is sensitive to stem-loop geometry. A) Secondary structure of *Ox40* wt and mirrored ADE RNA. Red nucleotides indicate point mutants required to stabilize the mirrored fold. $^{1}$H,$^{1}$H-NOESY spectrum of mirrored ADE imino region is shown on the right. The assignment is given in the 1D proton spectrum on top. Dashed lines indicate stretches with unknown directionality but known basepair identity. The confirmed basepairs are indicated by boxes in the secondary structure on the left. B) EMSA of coreROQ (left) and extROQ (right) binding to wt (top) and mirrored ADE (bottom). Protein concentrations are given on top. C) $K_D$s of EMSAs shown in B. D) $^{1}$H,$^{15}$N-HSQC spectra of apo core (top) and extROQ (bottom, black) and in presence of wt (left, magenta) and mirrored ADE (right, yellow). E) $^{1}$H,$^{15}$N-HSQC overlay of apo coreROQ (black) and in presence of wt (magenta) or mirrored (yellow) ADE.
